# New Evidence of the Earliest Domestic Dogs in the Americas

**DOI:** 10.1101/343574

**Authors:** Angela Perri, Chris Widga, Dennis Lawler, Terrance Martin, Thomas Loebel, Kenneth Farnsworth, Luci kohn, Brent Buenger

**Affiliations:** Department of Archaeology, Durham University, South Road, Durham DH1 3LE, United Kingdom; Don Sunquist Center of Excellence in Paleontology, East Tennessee State University, Johnson City, TN 37615, USA; Illinois State Museum Research and Collections Center, 1011 East Ash St., Springfield IL 62703, USA; Pacific Marine Mammal Center, 20612 Laguna Canyon Rd., Laguna Beach CA 92651, USA; Illinois State Museum Research and Collections Center, 1011 East Ash St., Springfield, IL62703-3500, USA; Illinois State Archaeological Survey, Prairie Research Institute, University of Illinois, Champaign, IL 61820, USA; Department of Biological Sciences, Southern Illinois University Edwardsville, Illinois 62026, USA; Western Archaeological Services, 1600 Dewar Dr., Rock Springs, WY 82901, USA

## Abstract

The domestication of dogs probably occurred in Eurasia by 16,000 years ago, with the initial peopling of the Americas potentially happening around the same time. Dogs were long thought to have accompanied the first migrations into the Americas, but conclusive evidence for Paleoindian dogs is lacking. The direct dating of two dogs from the Koster site (Greene Co., Illinois) and a newly-described dog from the Stilwell II site (Pike Co., Illinois) to between 10,190-9,630 cal BP represents the earliest evidence of domestic dogs in the Americas and individual dog burials in worldwide archaeological record. The over 4,500 year discrepancy between the timing of initial human migration into the Americas and the earliest evidence for domesticated dogs suggests either earlier dogs are going unseen or unidentified or dogs arrived later with a subsequent human migration.

---

The dog’s domestication and earliest uses have been topics of much debate in the archaeological and genomic literature, especially over the last decade (Germonpré et al. 2009; von Holdt et al. 2010; Larson et al. 2012; Germonpré et al. 2013; Thalmann et al. 2013; Freedman et al. 2014; Drake et al. 2015; Morey and Jeger 2015; Perri et al. 2015; Shipman 2015; Frantz et al. 2016; Perri 2016a; Morey and Jeger 2017). Advances in zooarchaeological, morphometric, and genomic methods have led to a burst of research in the field, but have also engendered disagreement regarding the interpretation of data from investigations of their origins. These debates extend to the earliest appearance of domesticated dogs in the Americas and the circumstances leading to their presence in the region, which are unresolved.

Though it is now widely accepted that all dogs were domesticated from ancient gray wolf ancestors (Vilà et al. 1997; Freedman et al. 2014), findings diverge on the timing, location, and number of domestication sites. The tentative identification of a number of proposed Paleolithic dogs dating from prior to the Last Glacial Maximum (Sablin and Khlopachev 2002; Germonpré et al. 2009; Ovodov et al. 2011; Germonpré et al. 2012; Germonpré et al. 2015a, 2015b; Germonpré et al. 2017), some up to 40,000 years ago (Camarós et al. 2016), has led to debate regarding the origins of the human-dog relationship (Crockford and Kuzmin 2012; Boudadi-Maligne and Escarguel 2014; Drake et al. 2015; Morey and Jeger 2015; Perri 2016a).

Despite the suggestion of domesticated dogs much earlier in the Paleolithic, a date of around 16,000 cal BP is generally accepted as the timing of domestication, based on secure archaeological and genomic evidence (Axelsson et al. 2013; Freedman et al. 2014; Morey and Jeger 2015; Frantz et al. 2016; Perri 2016a). Individual domestication locations have been proposed in the Middle East (von Holdt et al. 2010), Europe (Thalmann et al. 2013), Central Asia (Shannon et al. 2015), and East Asia (Wang et al. 2016), while Frantz et al. (2016) suggested a dual origin in both East Asia and Europe. The possibility of an independent domestication of dogs in the Americas has been raised by some (Koop et al. 2000; Witt et al. 2015), but rejected by others (Leonard et al. 2002; von Holdt et al. 2010).

The presence of early dogs in the pre-contact Americas is often assumed to be the result of companion animals arriving from across the Bering Land Bridge with migrating Pleistocene human populations (Schwartz 1997; Fiedel 2005; van Asch et al. 2013). Dogs may have assisted migrating groups by transporting goods and people, working as hunting aids, serving as bed-warmers, acting as alarms, warding off predators, and as a food and fur source. A recent analysis of dog remains from eastern Siberia suggests that dogs may have been important for hunting and particularly sled transport in the region up to 15,000 years ago (Pitulko and Kasparov 2017), similar to their present function in some Arctic regions today (Brown et al. 2013).

The earliest human migration into the Americas is proposed via a coastal route by between ~25,000 and 15,000 cal BP (Llamas et al. 2016; Skoglund and Reich 2016; Braje et al. 2017) or via a land route through the Ice Free Corridor by ~15,000 (Munyikwa et al. 2017; Potter et al. 2017). The earliest archaeological evidence of human presence in the Americas occurs in both North and South America around 14,500 cal BP (Dillehay et al. 2015; Halligan et al. 2016).

There are a number of large canid remains dating to the late Upper Pleistocene from across Beringia and southern Siberia, many of which are suggested to be Paleolithic dogs (see Germonpré et al. 2017 for a review of the Western Beringian and Siberian specimens). These include canids from Ulakhan Sular (c. 17,200 kya), Dyuktai Cave (c. 17,300-14,100 kya), Afontova Gora-1 (c. 16,900 kya), Verholenskaya Gora (c. 14,900 kya), Berelekh (c. 14,100 kya), Little John (c. 14,000 kya; Easton et al. 2011), McDonald Creek (c. 14,000-12,600 kya; Mueller et al. 2015), Nikita Lake (c. 13,800 kya), Ushki-I (c. 12,800 kya), and Ust’Khaita (c. 12,300 kya). At present, the taxonomy and interpretation of many of these specimens is contested or inconclusive. Others have yet to be evaluated further.

Although the arrival of domesticated dogs with an initial human migration has been the most reasonable explanation for their presence in the Americas, evidence for Paleoindian dogs has proven elusive. Previously, Jaguar Cave (Idaho) was thought to hold the earliest domestic dog remains in the Americas at over 10,000 years old (Lawrence 1967). However, when dated directly, the remains proved to be only 3,000-4,000 years old (Gowlett et al. 1987). Similarly, Beebe (1980) reported early dog remains dating to around 20,000 years ago from Old Crow Basin (Yukon Territory), but later dating demonstrated that this dog is of Late Holocene age (Harington 2003). While there are suggestions of domesticated dogs over 10,000 years old from a few North American sites (Haag 1970; Stanford 1978; Walker and Frison 1982; Grayson et al.1988; Saunders and Daeschler 1994; Jenkins et al. 2013; Lyman 2013), these canid remains have not benefitted from modern chronological or morphological (re)evaluation.

Fiedel (2005) suggested the lack of dog remains during the Paleoindian period is the result of their ephemeral presence, not their absence. While this is a distinct possibility, the earliest appearance of domestic dogs at Early Archaic sites in the midcontinent (Morey and Wiant 1992; Walker et al. 2005) raises questions regarding their origins and route into the Americas. Tito et al. (2011) reported finding the earliest evidence for dogs in the Americas at Hinds Cave (Texas) - a small bone fragment within a human coprolite. Genomic analysis was performed and the specimen was directly dated to around 9200 cal BP. Other early examples of domesticated dogs include specimens from Modoc Rock Shelter (c. 8,400 cal BP, Illinois; Ahler 1993), Dust Cave (c. 8,400 cal BP, Alabama; Walker et al. 2005), Rodgers Shelter (c. 8,800 cal BP, Missouri; McMillan 1970), and Koster (c. 9,500 cal BP, Illinois; Brown and Vierra 1983; redated in this study). Together, these specimens constitute the corpus of the earliest confirmed archaeological dog record in the Americas.

The arrival of dogs into the Americas has important cultural and ecological implications. Dogs were the first invasive species (along with humans) and domesticate in the Americas, potentially impacting populations of small mammals through predation, other species of *Canis* through hybridization, and other carnivores through transmission of diseases or competition (Doherty et al. 2017). They may have also contributed to important adaptations in hunting and mobility during the peopling of the Americas and into the Pleistocene-Holocene transition.

Here, we present the identification, analysis, and direct radiocarbon dating of an isolated dog burial from Stilwell II, an Early Archaic site in the Lower Illinois River Valley. We also present new direct radiocarbon dates for two dogs from the nearby Koster site. These dates confirm that the Stilwell II and Koster dogs represent the earliest directly-dated evidence for domesticated dogs in the Americas and the oldest intentional, individually-buried dogs known in the worldwide archaeological record. Other similar individual dog burials appear in hunter-gatherer contexts ~1,000 years later (Perri 2014; Perri 2016b). Importantly, we contribute to an emerging analytical framework for understanding the behavior and life history of these canids. Our analyses (zooarchaeology, paleopathology, morphology) lend insight into what these dogs looked like, how they lived, and their roles within Early Archaic communities.

## Site Backgrounds

### The Koster Site

The Koster site (11GE4) is located in a minor tributary valley of the lower Illinois River in Greene County, Illinois (Figure 1). The site is multicomponent and highly stratified, with cultural deposits spanning the Early Archaic to Mississippian, providing a nearly continuous record of Holocene human occupation (Brown and Vierra 1983). The site was excavated continuously over a ten-year period and is one of the most studied sites in the Lower Illinois River Valley (e.g., Butzer 1978; Hajic 1990; Komar and Buikstra 2003; Boon 2013).

**Figure 1.**
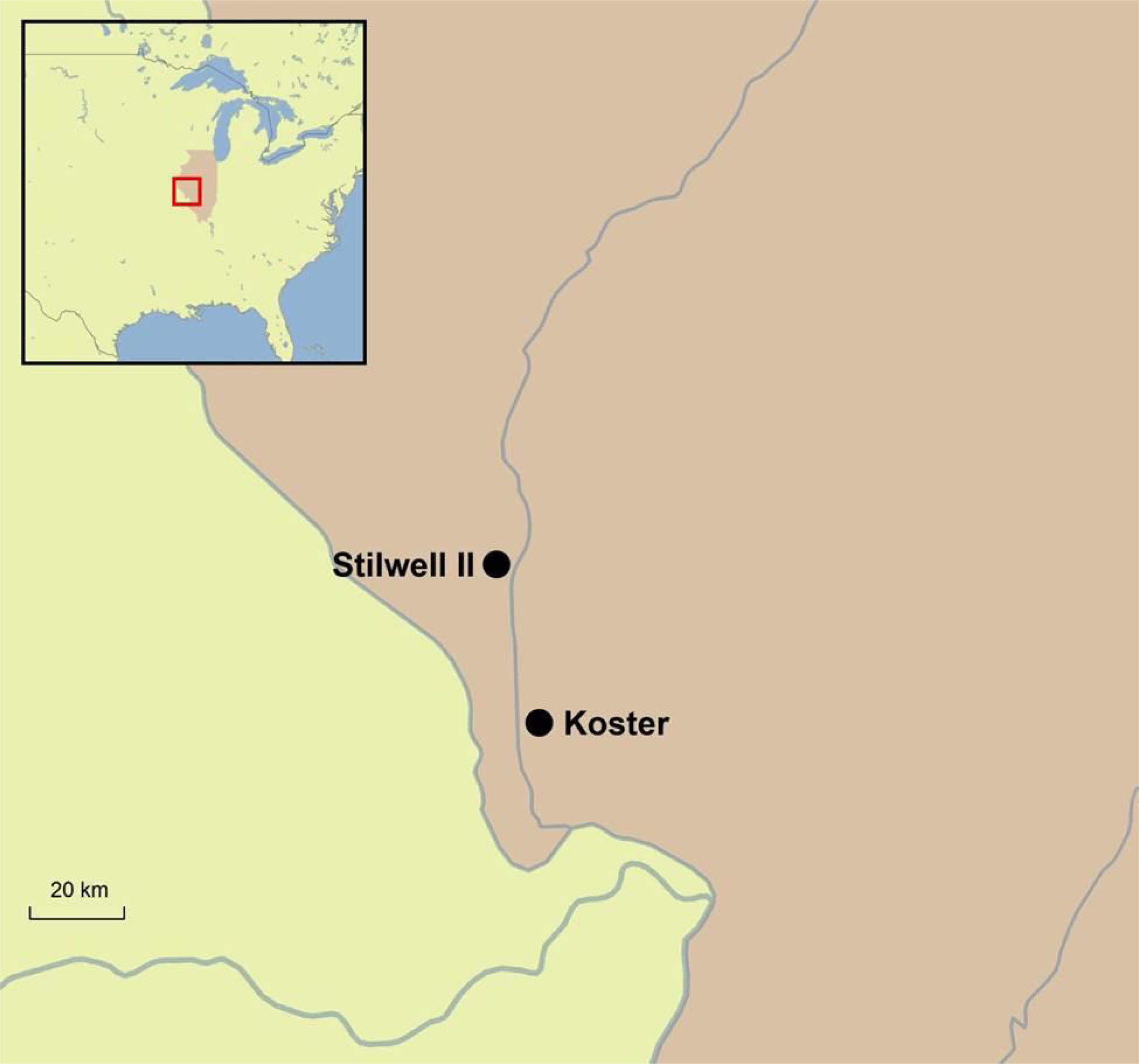
Location of the Koster and Stilwell II sites

Three isolated dog burials (cf. Perri 2017) in shallow, well-demarcated pits were identified from Horizon 11, one of the Early Archaic phases at Koster (Figure 2). There is also a fourth burial, likely associated with later period (Hill 1972; Morey and Wiant 1992), and a fifth burial is reported (Neusius 1996), the remains of which are not present in the current Illinois State Museum collection. The skeletons of the three Horizon 11 dogs were complete, articulated, and lacked evidence of butchering or skinning (Morey and Wiant 1992). Given their presence in Horizon 11 and association with a nearby charcoal date (Brown and Vierra 1983), the dogs were attributed to the terminal Early Archaic. Though this date is commonly reported as “8,500 years ago” (e.g., Morey and Wiant 1992: 225), the calibrated age based on the associated charcoal 14C dates is ~9,500 cal BP. These specimens are often cited as the earliest domesticated dogs and occurrence of intentional dog burials in the Americas (Morey and Wiant 1992; Fiedel 2005; Walker et al. 2005; Lapham 2010; Morey 2010).

**Figure 2.**
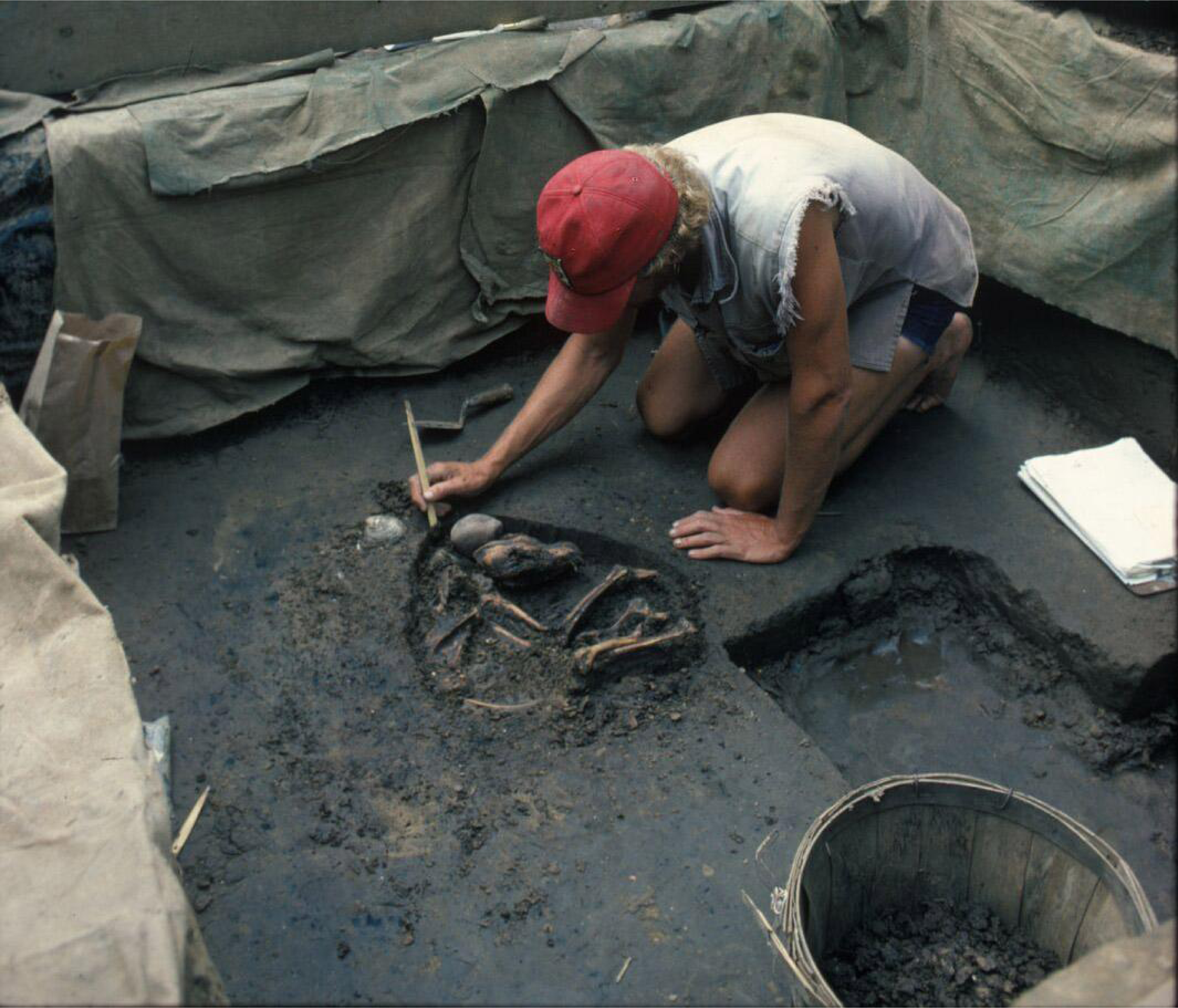
Excavation of the Koster F2256 dog burial (photograph by Del Baston, courtesy of the Center for American Archeology)

### The Stilwell II Site

The Stilwell II site (11PK1044) was discovered in 1960 when road grading operations cut through an alluvial fan in Pike County, Illinois, about 35km from the Koster site. Gregory Perino collected lithic and faunal remains from what he described as two living areas composed of a dark layer of soil, 6-inches thick and 20-feet long, exposed at the base of a 14-foot cut bank (Perino 1970:119). He subsequently recovered a dog burial in the northern area of the site, and a human burial in the southern area. The dog burial (Figure 3) was complete and articulated (Perino 1970, 1977). It was excavated and curated at the Illinois State Museum. The faunal remains collected by Perino and through later excavations by the Illinois State Archaeological Survey include white-tailed deer, turkey, turtle, small birds, vole, squirrel, fish and mussel shell. After the two rescue excavations in 1960 and 1962 Perino published very little about the site and left no field notes or maps. Re-excavation of the site began in 2015 by the Illinois State Archaeological Survey and is ongoing (see Supplementary Information).

**Figure 3.**
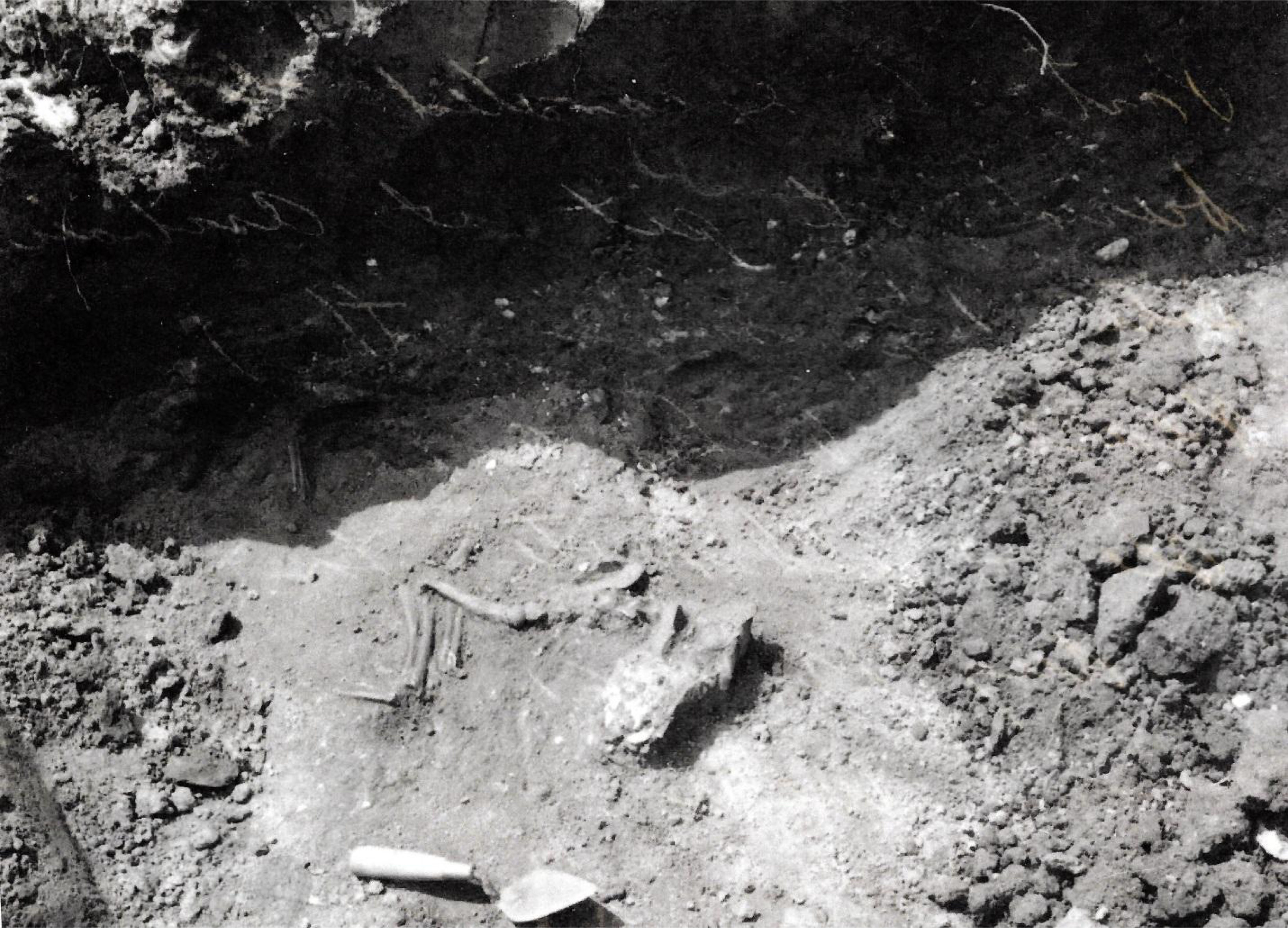
The Stilwell II dog burial *in situ* (photograph by W.L. Wadlow, courtesy of the Illinois State Museum Research and Collections Center)

## Materials and Methods

### Zooarchaeology, Morphology, and Paleopathology

The Koster and Stilwell II dogs were analyzed in the zooarchaeology laboratory at the Illinois State Museum’s Research and Collections Center, where they are curated. All skeletal specimens were examined in order to note the condition of epiphyseal closure, presence of cut marks, damage by carnivore or rodent gnawing, and exposure to fire. Analysis of shoulder height relies on the regression equations of Harcourt (1974), and body mass estimates use the methods presented by Losey et al. (2014, Losey 2016). Researchers have previously published comprehensive measurement data, burial information, and paleopathology for the Koster dogs (Morey 1992; Morey and Wiant 1992; Morey 2006; Lawler et al. 2016), which is not repeated here. Recent ancient DNA analysis of one Koster specimen has confirmed their status as domesticated dogs from Eurasia stock, likely originating in Siberia (Thalmann et al. 2013, Leathlobhair et al. 2018). Recovery of ancient DNA from the Stilwell II dog has failed thus far.

Though little documentation exists for the site, Perino (1970:119) is clear that the remains of the single Stilwell II dog were a shallow, intentional burial in what he described as the floor of a living area. The only *in situ* photograph of the dog shows an articulated skeleton as an isolated burial (Figure 3), with a northeast-southwest orientation, head facing east. The front legs appear to be tucked partly under the body.

Following von den Driesch (1976), we provide all possible skeletal measurements for the Stilwell II dog and have retaken all possible measurements from the two Koster dogs dated in this study (F2256 and F2357) (Supplementary Table 1). These measurements were compared to a sample of seven Archaic dogs from Iowa and Illinois (Supplementary Table 3). Modern wild canids *(C. latrans, C. lupus)* curated in the Illinois State Museum, the University of Kansas Biodiversity Institute, and the East Tennessee Museum of Natural History were also included in osteometric analyses to illustrate the morphological differences between domesticated and wild taxa. 3D models of the Koster F2256 and Stilwell II mandibles are available for download at www.morphosource.org (see Data Availability Statement).

Observations of the appendicular skeleton include overt and incipient pathology. We define incipient pathological changes as very mild or very early changes, not easily visualized by standard radiographic methods and not clarified substantially by standard computed tomography. Each bone was examined directly, supported by magnification as necessary. Microcomputed tomography has been conducted with some of the specimens, as parts of other studies (Lawler et al., 2016). All specimens were photographed. Observations were recorded by location within bone, thus resulting in multiple scores for given joint components (Supplementary Table 2).

### Radiocarbon Dating

Small rib fragments (1-2 cm in length) from the Koster and Stilwell II dogs were submitted to the University of Arizona AMS lab (Tucson, AZ) or Rafter Radiocarbon lab (Lower Hutt, New Zealand) for radiocarbon dating. In both cases, collagen was extracted using a modified Longin technique of acid demineralization followed by removal of organic contaminants using a weak basic solution (Longin 1971). Samples were combusted and further purified in a dedicated gas line and converted to graphite targets. These targets were analyzed using the accelerator at the Department of Physics, University of Arizona (USA) and National Isotope Centre, GNS Science (NZ), respectively. All 14C results are calibrated as 2-sigma age ranges with the Intcal13 dataset (Reimers et al. 2013) using Calib 7.1html (Stuiver et al. 2017).

## Results

### Zooarchaeology, Morphology, and Paleopathology

Aside from faint root etching on several of the long bones, examination of all Stilwell II specimens revealed only two occurrences of gnawing on the bones by small rodents. One is a small area (circa 6 × 3 mm) on the caudal surface of the right proximal humeral shaft. The second (circa 5 × 3 mm) is on the plantar surface of the left distal tibial shaft. No cut marks from dispatch wounds (e.g., on the atlas vertebra) or dismemberment (e.g., cuts near articular ends) are present on the skeleton. The dog is an adult of undetermined age and the absence of a baculum from the otherwise well-represented posterior skeleton suggests the animal was a female (see Supplementary Information).

Since it is a relatively complete skeleton, the Stilwell II dog has the potential to provide anatomical insights into the size and morphology of early North American dogs. It has an estimated shoulder height between 504-517 mm, based on radial (RDgl) and tibial (TAgl) length (Harcourt 1974) (Table 1). Losey et al., recently suggested improved methods for body mass estimation, based on cranio-dental (2014) and limb elements (2016). Application of these regression equations to the Stilwell II dog resulted in widely varying estimates (17-32 kg). Following Losey et al. (2016), we prefer body mass estimates that are based on elements relating directly to locomotion, such as limb elements. Estimates of body mass based on the humerus (distal breadth; HMbd) and radius (proximal breadth; RDbp) are both 17.1 kg (Table 1), similar in mass and build to a small modern English Setter. Contemporaneous dogs from the nearby Koster site are slightly shorter (shoulder heights: 439-463 mm) and more lightly built (12-14 kg) (Figure 4).

**Figure 4.**
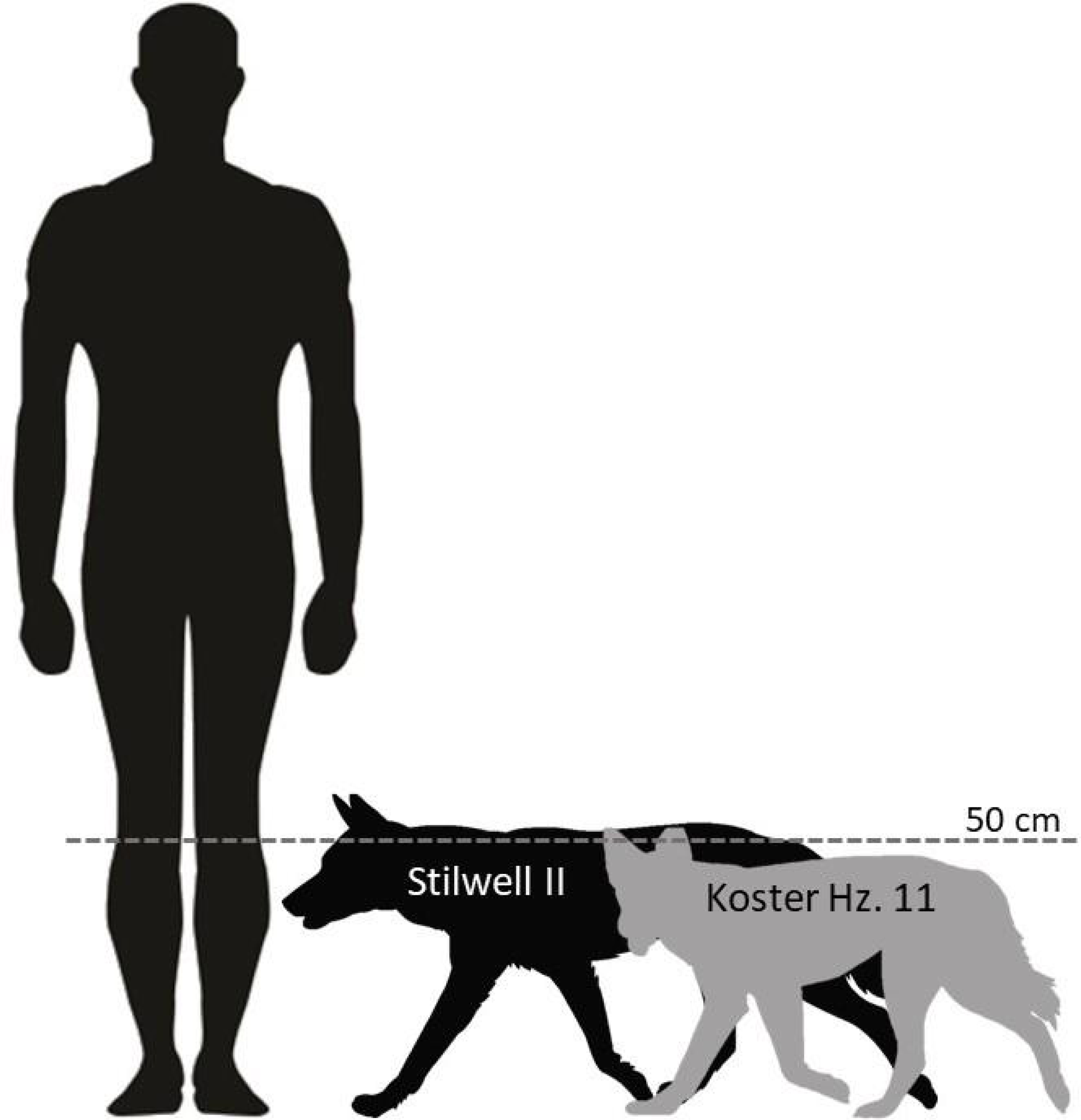
Relative size of the Stilwell II and Koster dogs.

**Table 1.** Shoulder height and body mass estimates of the Koster and Stilwell II dogs.

| Measurement | Shoulder Height (mm)<br>(Harcourt 1974) |  | Body Mass (kg)<br>(Losey et al., 2014) |  | Body Mass (kg)<br>(Losey et al., 2016) |  |
| --- | --- | --- | --- | --- | --- | --- |
|  | RDgl | TAgl | MR-18 | MR-19 | HMbd | RDbp |
| Koster F2256 | 450.7 | 439.8 | 13.4 | 15.3 | 12.5 | 13.6 |
| Koster F2357 | 463.1 | 453.0 | 16.5 | 16.9 | 12.8 | 14.3 |
| Stilwell II | 517.2 | 504.6 | 22.5 | 32.3 | 17.1 | 17.1 |

Mandibular morphology varies significantly between the Stilwell II and Koster dogs (Figure 5). The Stilwell II dog mandible is robust with relatively small carnassial molars and a deep mandibular body. Dog mandibles from the Koster site, however, are more gracile, with large carnassial molars and shallow bodies, relative to their size.

**Figure 5.**
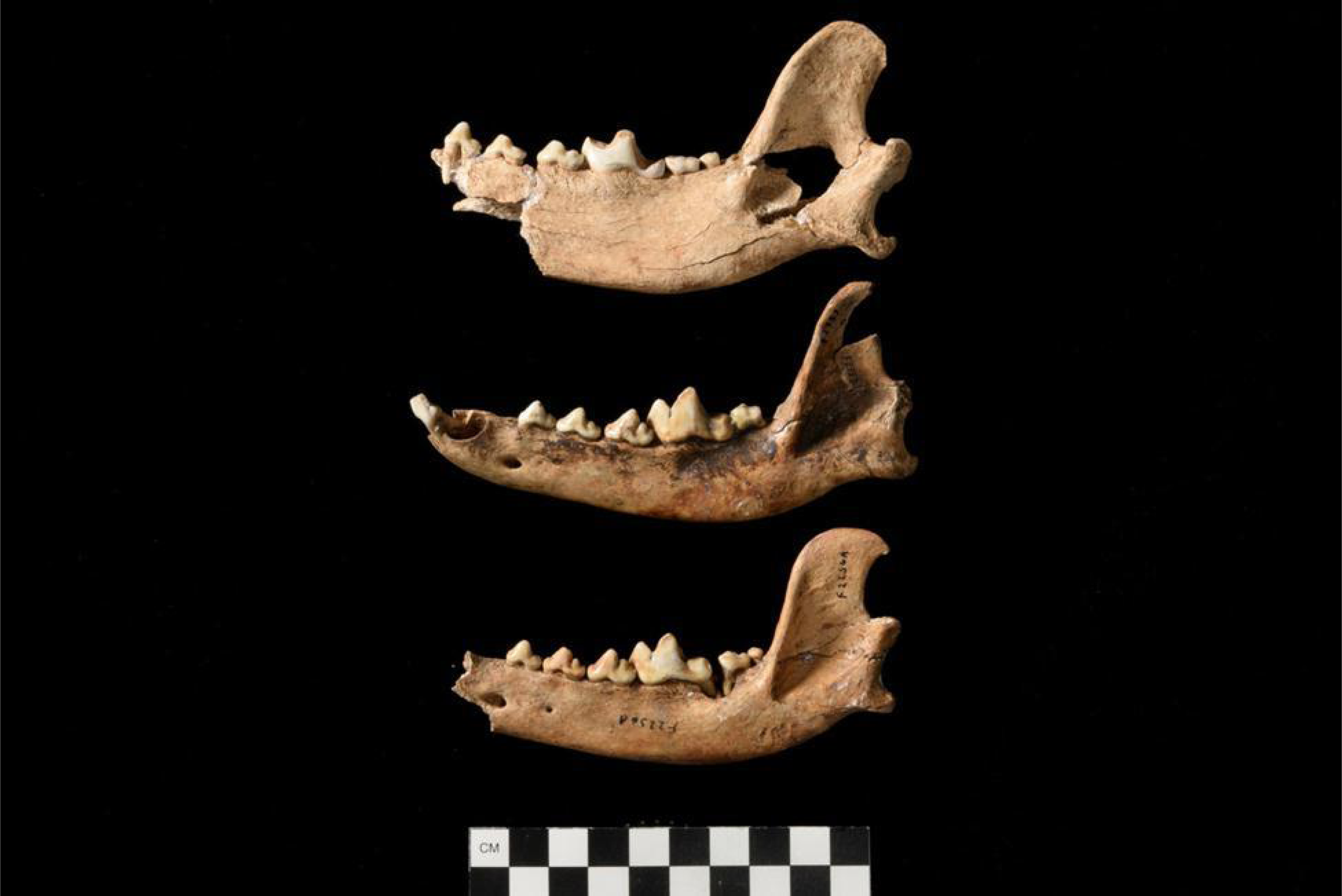
Comparison of left mandibles from the (top to bottom) Stilwell II, Koster F2357, and Koster F2256 dogs.

Observations of the Stilwell II dog’s axial skeleton included multifocal periodontal-periosteal disease and severe tooth wear. The first and second molars exhibit particularly extreme wear (Figure 6) and the right lower canine is worn nearly blunt. Damage of this type depends partly on genetic susceptibility, as in (e.g.) modern small breed dogs, and on diet and habits such as chewing on bones. The dog easily could have experienced several well-recognized complications of chronic oral cavity disease. The rough enlargement of perialveolar mandibular bone below the mandibular arcades signals gingival and periodontal disease. DeBowes and colleagues (1996) showed that multiple organ pathology can be related to oral cavity diseases such as gingivitis and periodontitis. In particular, significant associations were found between periodontitis and disease of the (a) kidney glomerularand interstitial tissue; (b) myocardium, especially papillary muscle; (c) hepatic parenchyma. The likely explanation is recurring bacteremia of oral tissue origin (Debowes et al. 1996). Without regular dental care, modern domestic dogs commonly develop similar oral pathology, and from the perspective of modern veterinary medicine, the Stilwell II dog would have been very uncomfortable.

**Figure 6.**
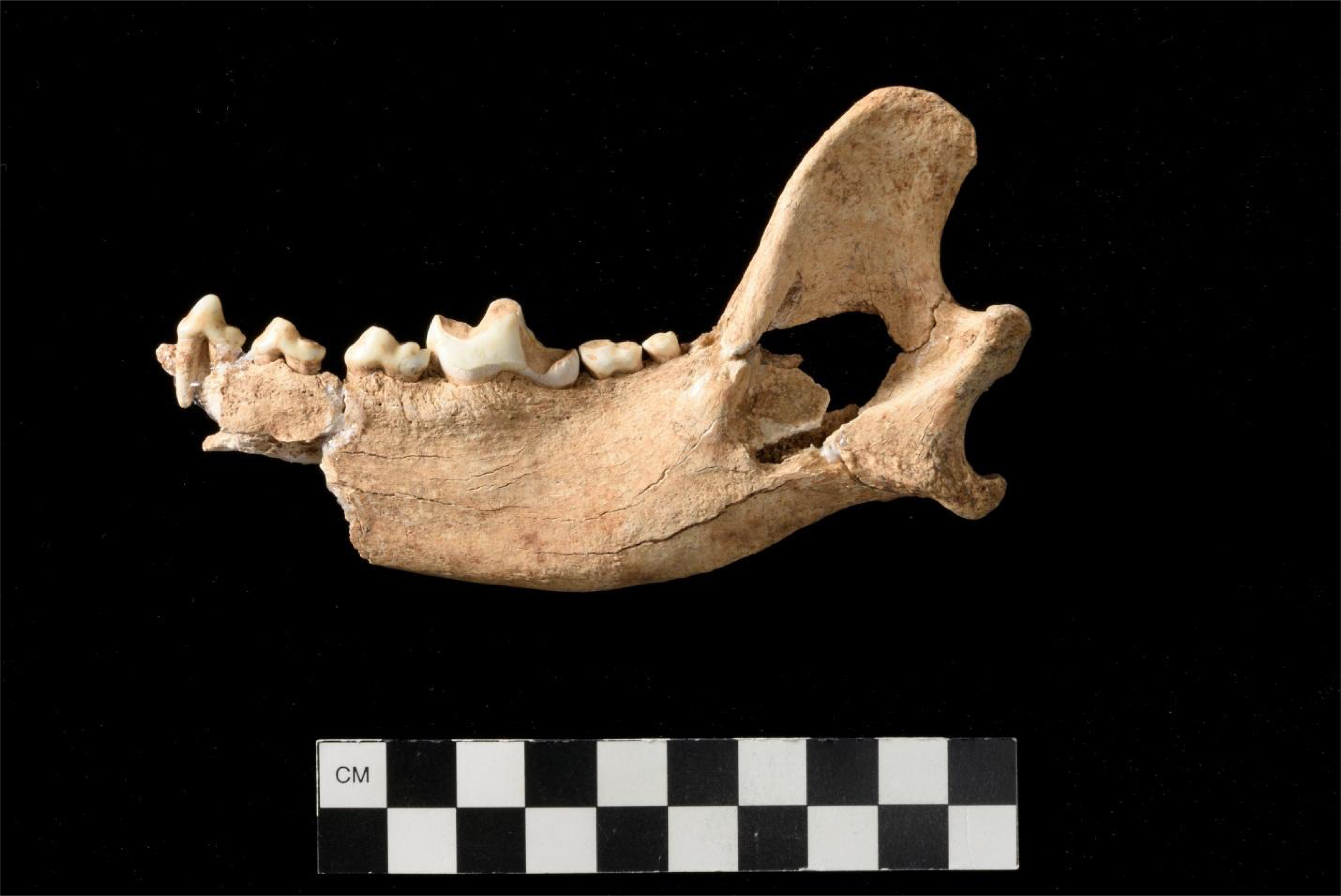
Left mandible of the Stilwell II dog showing advanced tooth wear.

Deviations of spinous processes were observed on seven vertebrae. Prevailing opinion has been that domestic dog vertebral spinous process deviations were caused by carrying packs or pulling travois (Darwent and Gililand 2001; Warren 2000; Walker et al. 2005). However, it has been shown recently that thorough differential diagnosis of these features yields multiple possible pathological causes or pseudopathologies (Lawler et al. 2016). Furthermore, the anatomical locations of affected vertebrae are protected by the caudodorsal neck ligament, tendon, and muscle mass (Miller et al. 1979) or lie below the protective transverse plane of the wings of the ilia. Thus, these vertebrae are not susceptible to injury related to carrying packs or pulling travois (Lawler et al. 2016) (Supplementary Table 2). A recent study of arctic foxes supports the notion that vertebral asymmetry can be a part of normal morphological variation in (at least) Canidae (Mustonen et al. 2017).

The limbs yielded observations of normal, incipient and overt pathological changes. The metapodials and phalanges yielded observations of incipient pathology (Supplementary Table 2). The summed changes are consistent with an active life style, and do not differ qualitatively from those that can be seen in modern adult dogs (Lawler and Evans 2016; Mustonen et al. 2017; Lawler et al 2017).

### Radiocarbon Dating

Neither the Stilwell II nor the Koster dogs have previously been directly radiocarbon dated. Based on their presence in Horizon 11, three dogs from Koster were associated with five Horizon 11 radiocarbon (^14^C) assays yielding dates between 8480 ± 110 BP (ISGS-236) and 8130 ± 90 BP (ISGS-1065) (Brown and Vierra 1983), but often cited as 8500 years ago (e.g., Morey and Wiant 1992). A fourth undated Koster dog likely comes from a later Archaic occupation. Here, we present three new direct ^14^C dates from the Stilwell II dog and two Koster Horizon 11 dogs (F2256 and F2357) (Table 2). Lyophilized samples from all three dogs had a white, fluffy appearance and carbon:nitrogen (C:N) ratios are within the range of modern mammalian collagen (2.9-3.6; Tuross et al., 1988), suggesting well-preserved collagen.

**Table 2.** Radiocarbon dates and stable isotope values for early dogs from the Lower Illinois River Valley.

| Lab No. | Specimen | Material | <sup>14</sup>C BP | 2-sigma, cal BP | C:N |
| --- | --- | --- | --- | --- | --- |
| AA101127 | Stilwell II dog | Bone collagen | 8840±80 | 10,190-9630 | 3.4 |
| NZA37136 | Koster Horizon 11 (dog F2256) | Bone collagen | 8790±30 | 10,110-9680 | 3.4 |
| NZA37135 | Koster Horizon 11 (dog F2357) | Bone collagen | 8820±30 | 10,130-9700 | 3.4 |

Koster dog F2256 dates to 8790 ± 30 BP (10,110-9,680 cal BP), Koster dog F2357 dates to 8820 ± 30 BP (10,130-9,700 cal BP), and the Stilwell II dog dates to 8840 ± 80 BP (10,190-9,630 cal BP). The chronological differences between the Stilwell II and Koster individuals are not statistically significant at the scale of 14C dating. These new dates range several hundred years earlier than previously-associated dates for the Koster dogs, and add another Lower Illinois River Valley dog to the early pre-contact dog record.

## Discussion

### Morphological Variation in Early North American Dogs

Most morphological work on North American dogs has focused on cranial shape (Morey and Wiant 1992; Morey 1992, Morey 2010; Olsen 1985; Walker et al. 2005); however, measurements on mandibles (Bozell 1988; Walker and Frison 1982) and limb elements (Morey and Aaris-Sorensen 2002) have also been examined. Although highly variable, North American dogs generally exhibit shortened muzzles with accompanying changes to dental and mandibular elements, relative to wild canids. Smaller body size, and the size of certain elements (i.e., carnassial molars) have been attributed to domestication (Morey 2010), though recent work has re-evaluated the usefulness of many so-called domestication markers (Janssens et al. 2016; Ameen et al. 2017; Drake et al. 2017). Unfortunately, crania are fragile and often poorly preserved in the zooarchaeological record. While relatively complete crania are present at the Koster site, the Stilwell II dog is represented only by cranial fragments despite field documentation indicating the presence of a complete skull.

To better understand morphological variability among early midwestern dogs, we use a limited set of mandibular measurements from a larger sample of Archaic midwestern domesticated dogs and modern wild *Canis* spp. (Supplementary Table 3). In this dataset, Archaic midwestern dogs generally have deeper mandibular bodies (i.e., greater height of the mandible behind the carnassial M1; von den Driesch 1976:60) relative to the length of the carnassial molar (von den Driesch 1976:60; Figure 7). Stilwell II, Simonsen, and one of the Modoc dogs have dog-sized carnassial molars, but relatively deep wolf-like mandibles. Three Koster and two Modoc dogs also have deep mandibles relative to carnassial size, although they are much smaller specimens. Coyote-dog hybrids (coy-dogs) generally plot near the archaeological dogs in this morphospace, suggesting that hybrid individuals may be difficult to distinguish on the basis of morphology alone. It is also possible that some of the early archaeological dog samples themselves are hybrid individuals, as suggested by recent ancient DNA analysis of one Koster dog (Leathlobhair et al. 2018).

**Figure 7.**
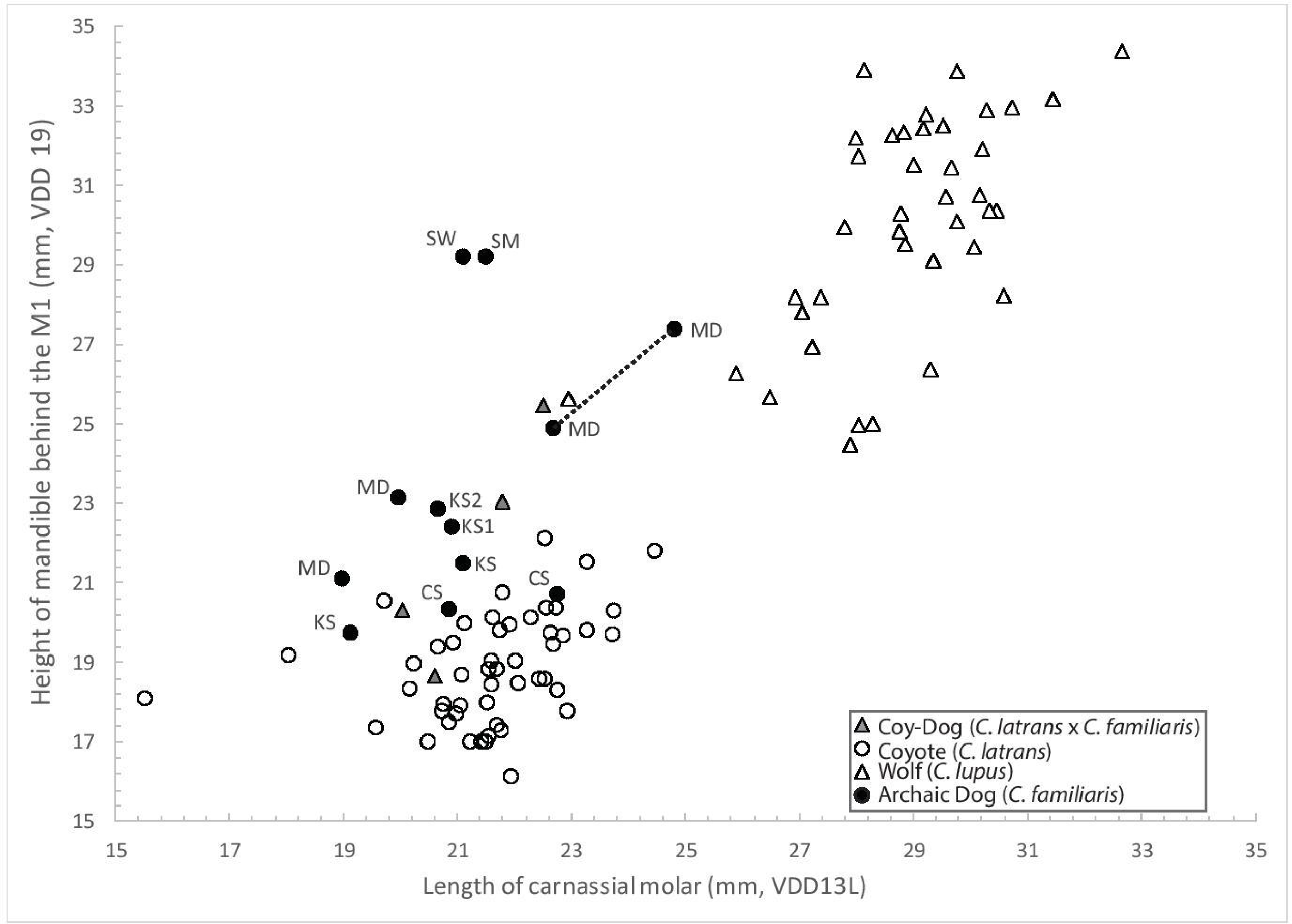
Relationship between carnassial length (von den Driesch 1976, measurement 13L) and mandibular body height (von den Driesch 1976, measurement 19) among different *Canis* groups. CS=Cherokee Sewer, IA; MD=Modoc Rock Shelter, IL; KS=Koster, IL (KS1 and 2 included in this study); SM=Simonsen, IA; SW=Stilwell II, IL. Dotted line connects left and right mandibles of the same individual. See Supplementary Table 3.

Even this limited sampling of Archaic dogs allows some comparative insight into early dog morphology, both contemporaneously and from several periods at the same site. For example, the three dogs from Modoc Rock Shelter show a significant range of variation in their mandibular height and length of carnassial molar (Figure 7). The largest dog dates to 8,560-8,200 cal BP (Supplementary Table 3), but exhibits intraindividual variation between the left and right mandible (shown via dotted line, Figure 7). Another dog from the site, dating to 5,710-5,330 cal BP, has much smaller molars and a more gracile mandible than that individual. A third undated, likely Archaic, dog falls between these two. Similarly, the two contemporaneous Koster dogs and a third undated, but likely contemporaneous, Koster dog all cluster together. A fourth dog, likely from a later period (Morey and Wiant 1992; Hill 1972), is smaller than those three in both measurements (Figure 7).

Though only a small sample, the distinct differences between the mandibles of the robust Stilwell II dog and the more gracile Koster dogs (Figure 5), from individuals geographically and temporally indistinguishable, suggest there may have been some amount of variation already in the earliest American dogs. This is perhaps unsurprising given the morphological variation seen at sites from similar time periods in eastern Siberia (Pitulko and Kasparov 2017). The probable female sex of the larger Stilwell II dog suggests the morphological differences are not the result of sexual dimorphism, especially given the similarly gracile directly-dated Koster dogs are both a female (F2256) and a male (F2357) (Morey and Wiant 1992). Similar morphological variation is also seen in the two contemporaneous Middle Archaic dogs from Iowa (Figure 7), with one being more robust like the Stilwell II dog and the other more gracile like the Koster dogs. Though this variation may be the result of morphologically distinct American dog lineages, it may also arise from local admixture with wild canids, such as coyotes and wolves, leading to rapid variation within a more homogenous initial dog population.

### Hybridization of Early North American Dogs

Genomic work on wild canid populations has demonstrated that all North American *Canis* spp. have the ability to interbreed, often to a significant degree (Wayne and Jenks 1991; Monzón 2014). Although we analyze these taxa as distinct groups, it is likely that some of these specimens show admixture of different species, even in groups made up of modern museum specimens of “known” taxonomic affinity. For example, a recent ancient DNA analysis of one of the Koster dogs we dated (F2256) found that although the specimen clusters with all other pre-contact North American dog material analyzed (spanning c. 9,000 years), it also shows evidence for potential admixture with a midwestern coyote (Leathlobhair et al. 2018). This may account for the morphological variation seen between the Koster dogs and the Stilwell II dog, which are otherwise concurrent in space and time.

For these reasons, we consider the present study as merely illustrative of general morphological trends in archaeological *Canis.* Combined genomic and morphological approaches have the potential to answer many lingering questions about North American dog populations. However, these techniques are just beginning to be applied rigorously to questions of early dogs in the Americas.

## Conclusions

The c. 10,000 year-old age of the Stilwell II and Koster dogs introduces an over year gap between these earliest dog remains and the proposed initial migration into the Americas. This is consistent with genomic analyses suggesting North American dog populations originated around 10,000 cal BP (Witt et al. 2015), though others suggest an earlier split from Eurasian dogs (Thalmann et al. 2013; Leathlobhair et al. 2018). It is possible that the arrival of the first human populations into the Americas predated their access to Eurasian domesticated dogs and thus they arrived without them. In this scenario, dogs may have arrived with later migrating Siberian groups (as part of ongoing “migratory dribbles”; Meltzer 2009: 200) via an interior route before 10,000 cal BP, but were not part of the very first pulse of migration into the Americas.

The Bering Land Bridge was flooded by ~11,000 years ago (Jakobsson et al. 2017), but populations may have crossed prior to this period, moving into North America through the Ice Free Corridor and dispersing into the midcontinent, leading to the earliest dog records in this region beginning at ~10,000 years ago. This scenario would posit that earlier domestic dog remains have yet to be found in Alaska, western Canada, the northern Great Plains, the Intermountain West and dispersed near the southern exit of the Ice Free Corridor. Many potential early dog remains fit with these circumstances, including those from McDonald Creek (Alaska; Mueller et al. 2015), Little John (Canada; Easton et al. 2011), Agate Basin (Wyoming; Walker and Frison 1982), Danger Cave (Utah; Grayson 1988), and Hogup Cave (Utah; Haag 1970). Equally, undated canid remains from the Early Archaic layers of Daisy Cave on California’s Channel Islands may represent the earliest dog remains from the western coast (Rick et al. 2008).

The earliest New World domesticated dogs appearing in the midcontinent around 10,000 years ago present a conundrum both temporally and spatially, but the current absence of Paleoindian dogs in the west may be the result of several factors. Earlier dogs in western North America may be going unseen or unrecognized, despite the plethora of Clovis, WST and earlier sites (Erlandson et al. 2011; Smallwood and Jennings 2014; Stanford and Stenger 2014). Some regions still have few early sites and the ephemeral nature of some sites (e.g., procurement or satellite camps) may constrain the discovery of dog remains (Fiedel 2005; Erlandson et al. 2011). If early dog remains are being encountered, they may not be identified as dogs, particularly given the often limited and poorly-preserved nature of early skeletal material and difficulties in distinguishing early dogs from wolves (Perri 2016a) and coyotes. The few specimens that have been tentatively proposed as Paleoindian dogs (e.g., Haag 1970; Walker and Frison 1982; Grayson 1988) have not been re-evaluated, leaving their taxonomy unclear. These potential dogs also date to not much earlier than the Stilwell II and Koster specimens, leaving any dogs associated with the earliest human migration in the Americas unaccounted for. Finally, it is possible that domesticated dogs entering the Americas with human groups facilitated rapid movement into the midcontinent, leaving little trace in western North America.

The foregoing observations raise the question of whether domesticated dogs ever accompanied humans across Beringia. While an *in situ* domestication of North American wolves has been raised as a possibility (Koop et al. 2000; Witt et al. 2015), this has been rejected by several genetic analyses (Vilà et al. 1997; Leonard et al. 2002; von Holdt et al. 2010; Freedman et al. 2014; Leathlobhair et al. 2018). Some North American archaeological dog specimens do show genetic similarities with North American wolves (Koop et al. 2000; Witt et al. 2015), however this is likely the result of admixture rather than North American wolf domestication. Additional work on ancient American canids, particularly the inclusion of more ancient North American wolf and coyote reference specimens, will further clarify this issue.

Identification of earlier Paleoindian dogs, if they exist, will require distinguishing them from wild canid taxa. This has proven a difficult task, as seen from debates regarding proposed early dogs in the Paleolithic record of Eurasia (Ovodov et al. 2011; Crockford and Kuzmin 2012; Boudadi-Maligne and Escarguel 2014; Drake et al. 2015; Germonpré et al. 2015; Perri 2016a). Differentiation between wild and domestic canids has been based primarily on morphological traits, often requiring well-preserved cranio-dental material. The validity of these traits for distinguishing domestication is also questionable, given the morphological plasticity of *Canis* (Morey and Jeger 2015; Janssens et al. 2016; Ameen et al. 2017; Drake et al. 2017). Substantial introgression between newly-arriving Eurasian dogs and North American wolves and coyotes likely contributed significantly to American dog ancestry as well (Leathlobhair et al. 2018). This potentially extensive introgression, particularly in the case of early coy-dogs (dog x coyote hybrids), may contribute to the misidentification of these specimens in the archaeological record. Though some past research has emphasized apparent introgression between ancient dogs and coyotes or wolves (Walker and Frison 1982; Valadez et al. 2006), the issue of early hybridization warrants more attention in future studies.

Analysis of ancient DNA is increasingly being used to identify domesticated dogs (Larson et al. 2012; Druzhkova et al. 2013; Frantz et al. 2016), but requires adequate preservation of skeletal material and is subject to debates about the interpretation of results (Savolainen et al. 2002; Ding et al. 2012; Thalmann et al. 2013; Skoglund et al. 2015). Increasingly, techniques that do not rely on ancient DNA preservation or preservation of pristine specimens, such as complete crania, are allowing researchers to document individual life histories of canids, improving chances of identifying individuals in close contact with humans. These techniques include investigating paleopathology and trauma to clarify, for example, pack-loading and mistreatment (Losey et al. 2014; Lawler et al. 2016) and geometric morphometrics (GM) to detect biomechanical differences among canids (Drake et al. 2015; Evin et al. 2016; Drake et al. 2017). Dietary analysis of stable isotopes may also help to identify early canids in close contact with humans (Ewersen et al. 2018). Ultimately, a combination of these methods will best promote the identification of the earliest domesticated dogs (and other domesticated species).

The Stilwell II and Koster dogs were contemporaneous adult, medium-sized dogs with very active lifestyles and varied morphologies for their proximity in space and time. Early American dogs likely played key cultural and ecological roles in the movement and adaptation of migrating human populations and their intentional burial suggests they were an important part of human domesticity in the Americas by the late Paleoindian/early Archaic. Similar intentional dog burials in other temperate hunter-gatherer contexts have been associated with the dog’s importance as adaptive tool technology in the face of changing environments and prey during the Pleistocene/Holocene Transition and their subsequent elevated social status (Perri 2014; Perri 2016b). The intentional burial of the Koster and Stilwell II dogs may reflect a similar importance of hunting dogs in the deciduous forested environment of the midcontinent.

The dating of the Stilwell II dog to around 10,000 years ago, coinciding with similar dates for the Koster dogs, adds a further early specimen to the pre-contact dog record and identifies the Lower Illinois River Valley as a site of early North American domestic dog activity. These new dates extend the presence of North American dogs potentially into the Dalton period, a transitional Late Paleoindian-Early Archaic phase in the Midcontinent (Koldehoff and Loebel 2009). They also confirm the Stilwell II and Koster specimens as the earliest dogs in the Americas and earliest examples of intentional, individual dog burials in the worldwide archaeological record^1^. Future (re)evaluation of faunal remains from Clovis, Western Stemmed and earlier sites may further identify domesticated dogs in the earlier Paleoindian record, supporting their arrival with the first human migrations into the Americas. Alternatively, the timing and location of these earliest dogs may suggest a later arrival with subsequent early human migrations.

## Footnotes

1 Although an individual dog burial has been reported from the Siberian Beringian site of Ushki-1 (Dikov 1979), which dates to around 13,000 years ago (Goebel et al. 2010), by all accounts these remains were identified via photograph in the 1970s and are now lost (Pitulko and Kasparov 2017), having never been confirmed as a dog or directly dated.

## Acknowledgements

We thank the Illinois State Museum for access to the faunal collections, the Center for American Archeology for use of the Koster excavation photograph, and Doug Carr and Claire Martin for additional material photographs. We also thank Bonnie Styles, Dee Ann Watt, Mike Wiant, Beckie Dyer, Mike Kolb and Jim Theler for their assistance and Ken Ames, Torben Rick, and anonymous reviewers for their helpful comments on the manuscript. ISAS would particularly like to acknowledge Bob Perino for the generous donation of his father’s material from the Stilwell II site to them. Funding was provided by the Illinois State Museum Society, the ETSU Center of Excellence in Paleontology, the Rosemary Cramp Travel Fund and the R. Bruce McMillan Museum Research Internship (A.R.P).

## Data Availability Statement

3D models of the Koster F2256 (http://www.morphosource.org/Detail/SpecimenDetail/Show/specimenid/10494) and Stilwell II dog (http://www.morphosource.org/Detail/SpecimenDetail/Show/specimenid/10495) mandibles are freely available on Morphosource.

## Supplemental Materials

For supplementary material accompanying this paper, visit www.iournals.cambridge.org/Mournal].

### The Stilwell II Site

In 1962, Perino returned to the Stilwell II site and conducted further exploratory excavations, at which time he recovered additional material consisting of broken projectile points, chipped stone tools, lithic debris, and faunal remains. Perino suggested that the site dated to “approximately Dalton times” (1977:99) based on the tool technology. Two of the broken projectile points he recovered were used as type specimens for the Stilwell variant of Kirk corner-notched points (Perino 1970). Perino reported that a bifurcate base point similar to LeCroy or Fox Valley, a Dalton adze, and the base of an Agate Basin point were also recovered during this second dig (Perino 1970, Perino 1977). Perino estimated the Fox Valley and Stilwell points dated to ~7000 BC (1985:136, 365), and the Agate Basin points to ~8300-8000 BC (1985:5), suggesting that more than one occupation component (~1000 years apart) may be associated with the Stilwell II Early Archaic deposits. Agate Basin points are now known to date to between ca. 12,100-11,800 cal BP (Pelton et al. 2017), earlier than Perino’s estimate.

Perino’s observations introduced the possibility that the human burial and dog burial may not be contemporaneous features. Despite Perino’s opinion of the importance of the site, he conducted no further work there and only minor mentions of the site appeared subsequently. No field notes or maps exist, aside from a drawing of the human burial (Farnsworth 2006: 66) and an *in situ* photo of the dog burial (Figure 3). A series of black and white prints and color slides taken during the 1960 roadwork are curated at the Illinois State Museum Research and Collections Center.

The extent of the additional work conducted by Perino in 1962 is unknown, but based on the black and white photos, is assumed to have been focused along the western cut face. Perino also suspected that the site deposits extended back under the alluvial fan and beneath the road. Perino retained possession of the material from his 1960 and 1962 work, with the exception of the dog and human burials, and some scattered finds of Archaic and Woodland period points from the general area. These were given to Pat Munson who transferred them to the Illinois State Museum.

In 2014, the remaining non-burial material from the site was donated to the Illinois State Archaeological Survey (ISAS) by Perino’s son. The donated assemblage contained numerous well-preserved faunal remains, lithic debris, and the broken points and chipped stone tools from the site. These appeared previously in Perino’s short published comments on his site work at Stilwell II (Perino 1962, Perino 1977), confirming the origins of the collection.

Included in the lithic assemblage were 309 lithic artifacts plus faunal remains. Lithic debitage (*n* = 271) comprised the bulk of material, dominated by large (1” - 1½” size class) core reduction flakes. Twenty-seven chipped stone tools were also present, including the proximal fragments of two Stilwell points, five biface/preform fragments, one drill/perforator fragment, nine utilized/edge damaged flakes, two chipped stone adzes (with wood use wear polish), one adze preform, six multidirectional cores, and one lanceolate point proximal fragment. Two hammerstones and two faceted groundstone cobbles, one of which is stained with red ochre, complete the assemblage. Seven miscellaneous items were also present, including one fragment of fire-cracked rock (FCR), a coral fossil, a pebble, and three abraded limonite and hematite fragments.

The assemblage is biased toward larger-sized artifacts, likely reflecting a selective collection strategy by Perino. Thus, while it is difficult to interpret the lithic assemblage, it is clear that occupants of the site were engaged in the procurement and primary reduction of the abundant outcrops of Burlington chert that surround the site. Five artifacts attributed previously to the Stilwell II site in published photographs are extant in the donated assemblage (Perino 1962:48). Other diagnostic points in the collection include the proximal fragment of a lanceolate point resembling an Agate Basin point. The bifurcated base point mentioned by Perino in his 1970 publication is not present.

Faunal remains recovered by Perino include a wide variety of species such as white-tailed deer, small birds, turkey, turtle, fish, and mussel shell fragments that are typical for the Archaic period in this region. A few of the larger specimens show obvious cut marks from butchering, and several modified bone tools are also present.

In 2015, personnel from ISAS visited the Stilwell II location in an effort to relocate the site and evaluate the potential for remaining intact deposits. A topographic map of the site area was generated and 13 bucket augers were excavated along both sides of the road cut. All sediments were screened through 1/4” mesh, with the exception of one auger that was retained for flotation processing. Most augers were positive and indicated the presence of lithics, faunal remains, burned earth, and charcoal at depths up to 85 cmbs (limits of testing) along the base of the western cut face. Numerous pieces of debitage (*n* = 92), one perforator fragment, and two utilized flakes were recovered from depths of 20-85 cm during auger testing. Recovered faunal material included 1015 pieces, primarily from the nine-liter flotation sample. Although most of these were too small to identify to species, the assemblage consists primarily of fish (bowfin, catfish, gar) and squirrels, with some bones being burned. Fragments of vole, eastern cottontail, raccoon, and a possible elk dew claw were identified as well.

Five 2-inch Giddings probe cores were placed across the longitudinal profile of the fan which identified buried paleosols, including anthropogenic deposits, on top of the fan at a depth of 1.5 m, and within an ACb horizon along the roadside at depths of 0.16 m to 1 m (Kolb 2015). A Bw horizon containing artifacts at depths up to 1.5 m is also present, suggesting an even older component at the site. East of the road, the distal fan lies within an active agricultural field with artifacts of varying ages (Archaic-Woodland) exposed on its surface and incorporated into the plow zone. The artifacts from the 2015 work are currently being analyzed, and investigations at the site are ongoing.

### The Stilwell II Dog

At or since excavation, transport, and storage, some of the elements which appear intact in the *in situ* 1960 photograph, such as the skull, have collapsed. Inspection of the skeletal material revealed approximately 200 specimens (weighing a total of 678.5 g), 177 of which are identifiable to one individual adult domestic dog. These include 14 cranial fragments, the posterior portions of both mandibles, and 3 isolated teeth; 17 whole and fragmented long bones having closed epiphyses (proximal ends of both humeri and both femurs are missing); 32 vertebral fragments; 55 rib fragments; 47 foot bones (i.e., carpals, tarsals, metapodials, and phalanges); 5 small fragments of scapulae; and 2 incomplete pelves (Supplementary Table 1). A baculum is not present in the inventory, although the posterior skeleton is otherwise well-represented. Therefore, the tentative sex of this individual is female or indeterminate. The Stilwell II dog is curated at the Illinois State Museum Research and Collections Center.

Six other specimens were found mixed among the dog bones. These include a left pelvis fragment (ilium and acetabulum) from a female white-tailed deer *(Odocoileus virginianus);* left tibial fragments from two gray squirrels *(Sciurus carolinensis);* a left ulna shaft from a medium-sized duck (Subfamily Anatinae); a right femur from a passenger pigeon *(Ectopistes migratorius);* and an anterior carapace fragment (part of a proneural bone) that is most comparable to painted turtle *(Chrysemys* picta). These are likely habitation debris that was included unintentionally with the burial fill.

**Supplementary Table 1.**
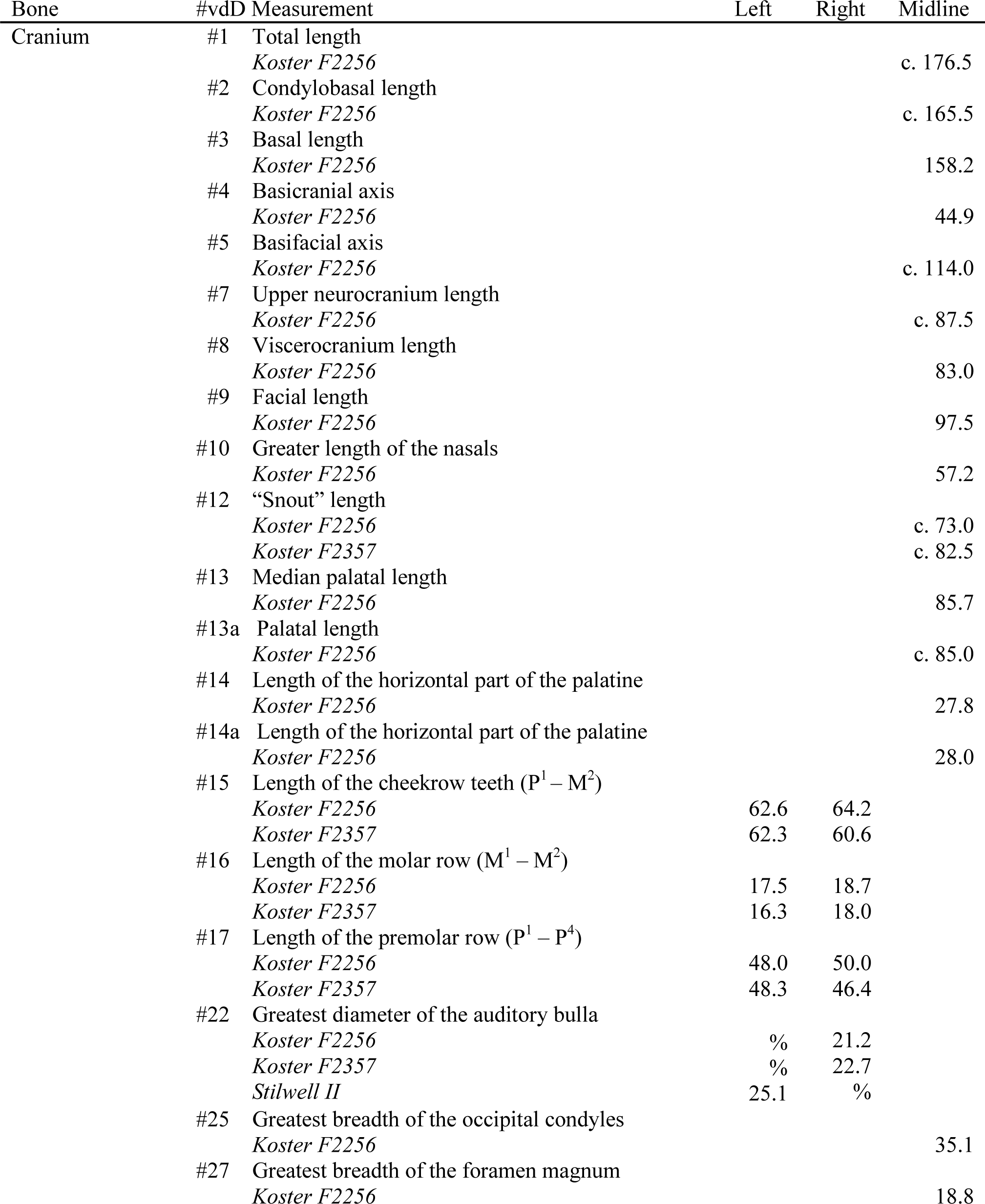
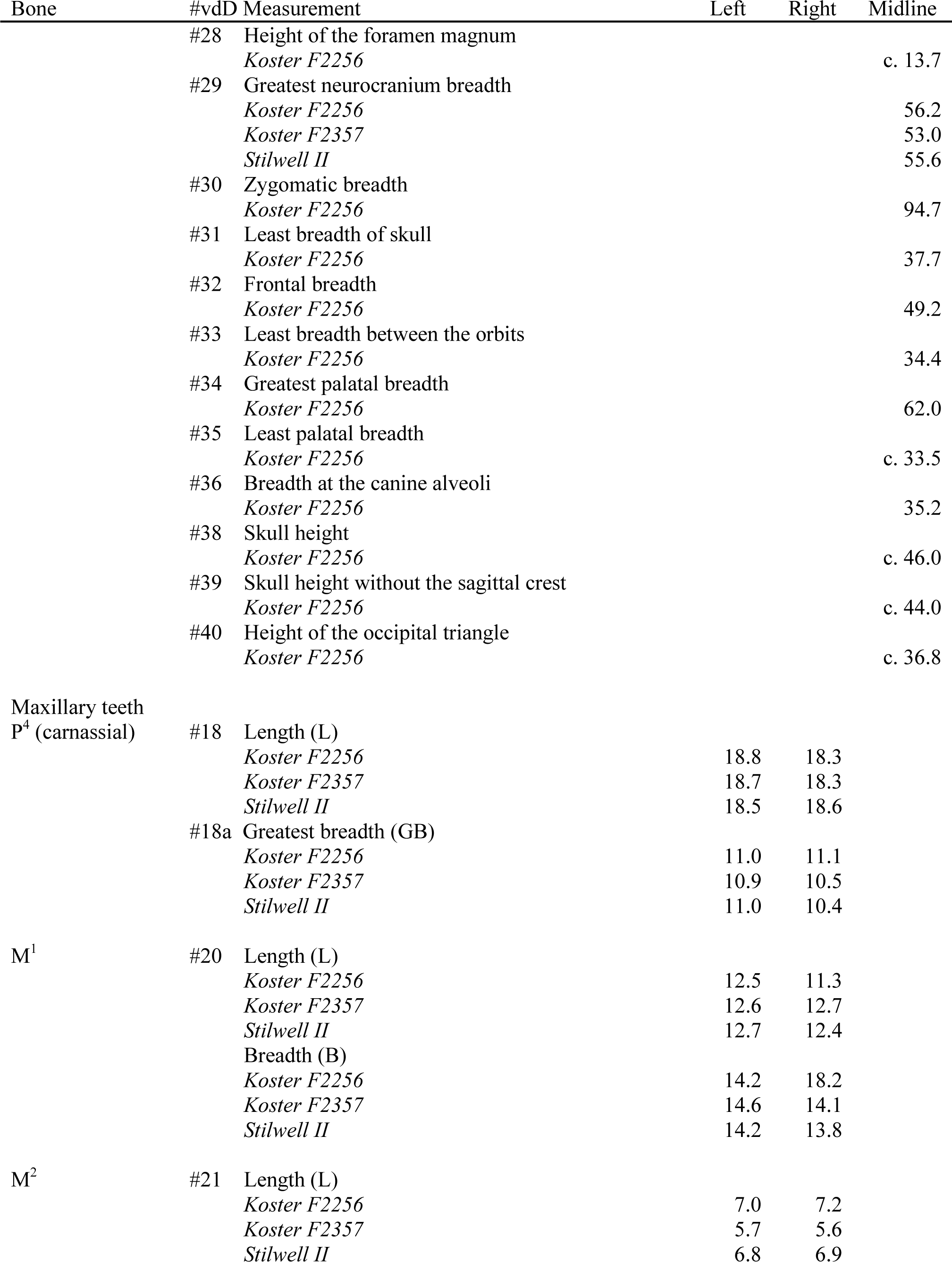
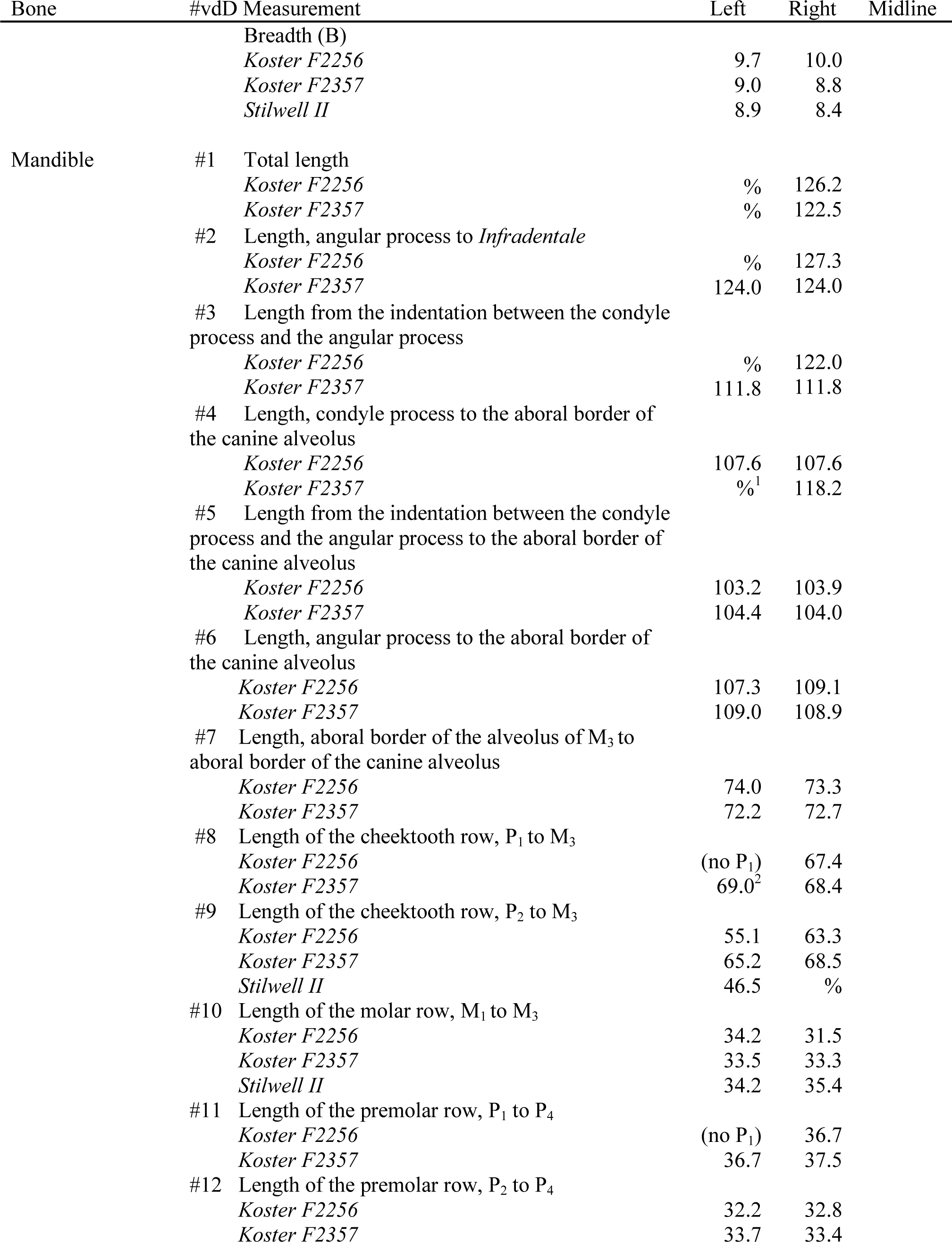
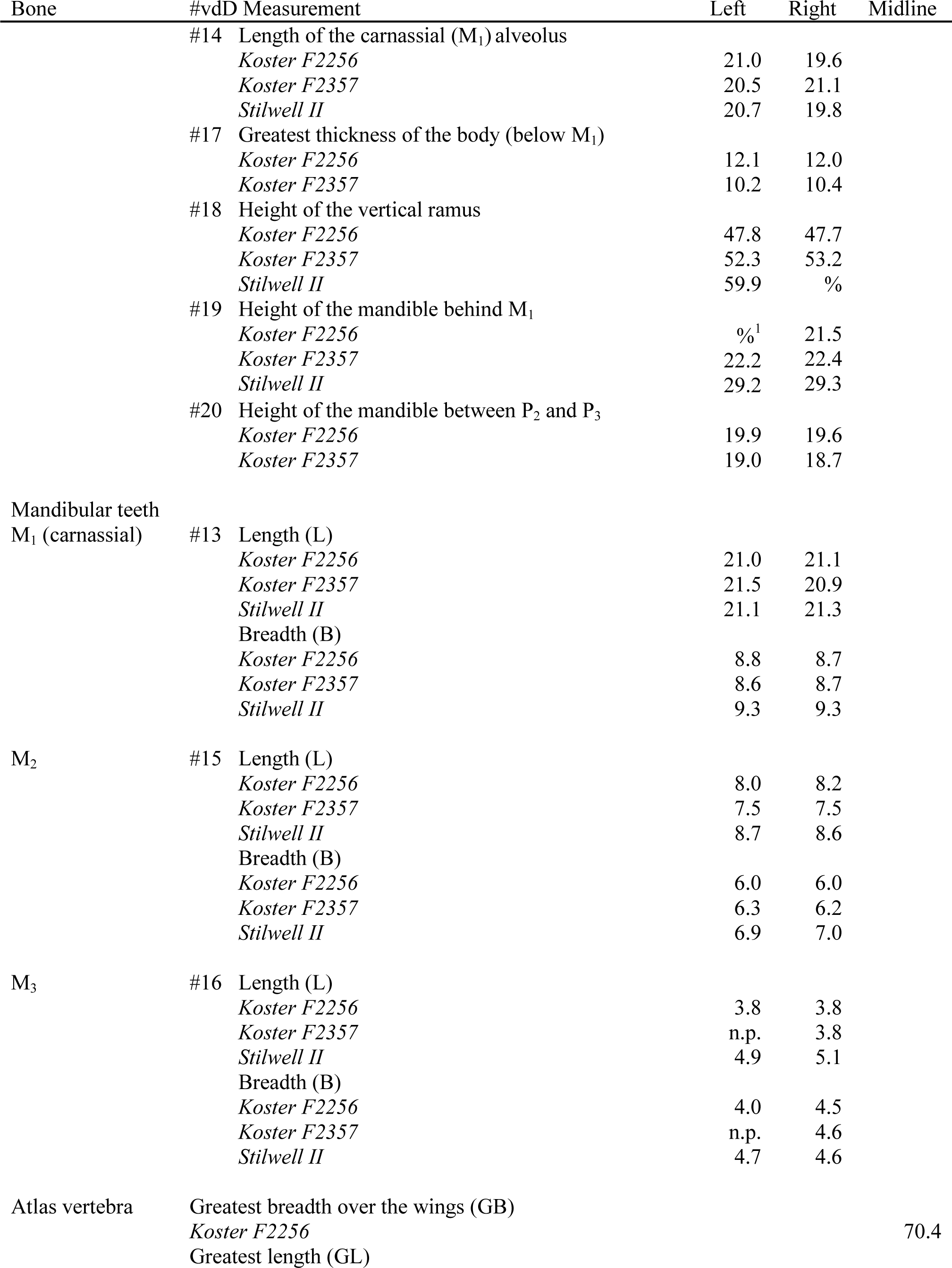
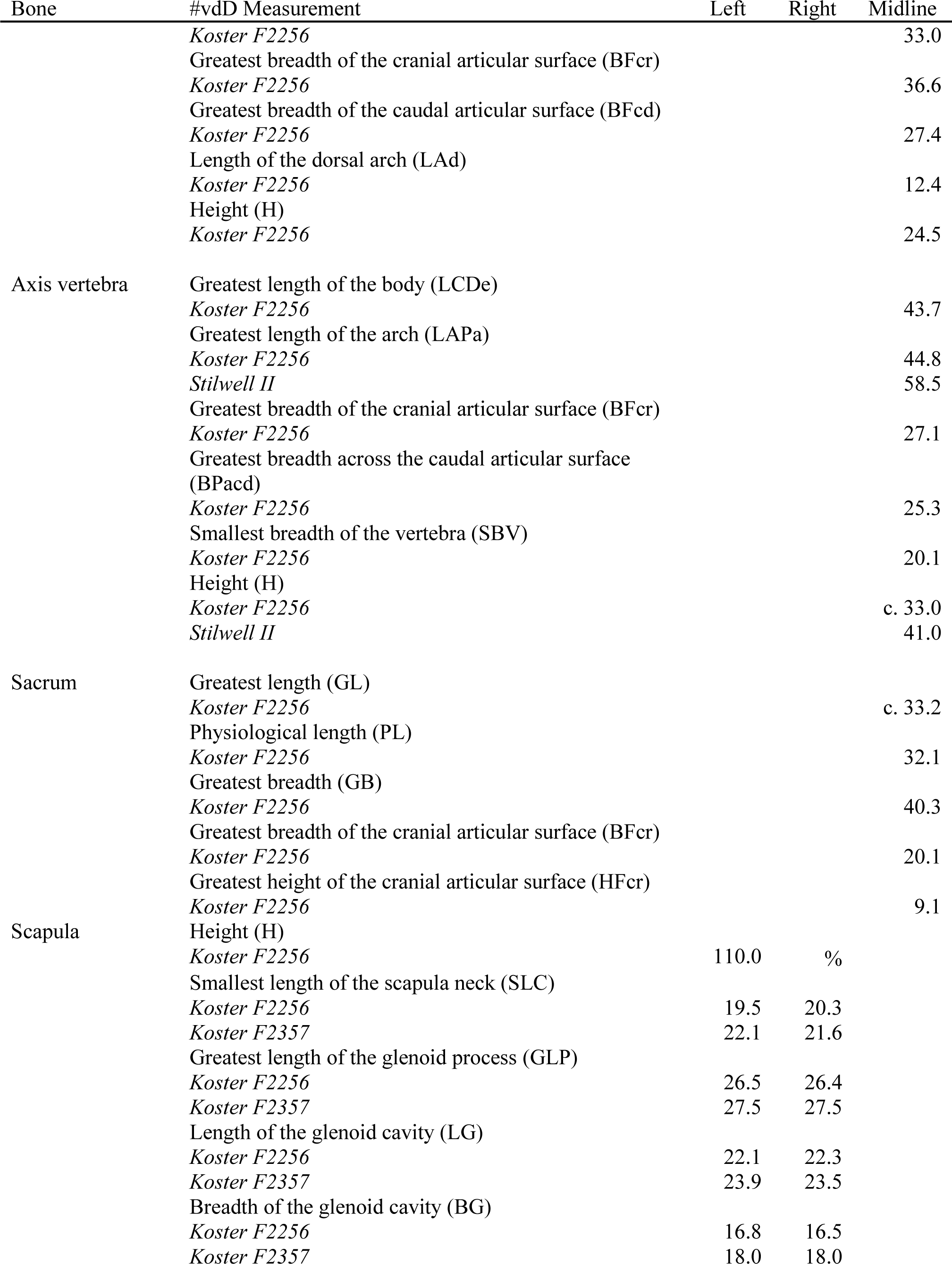
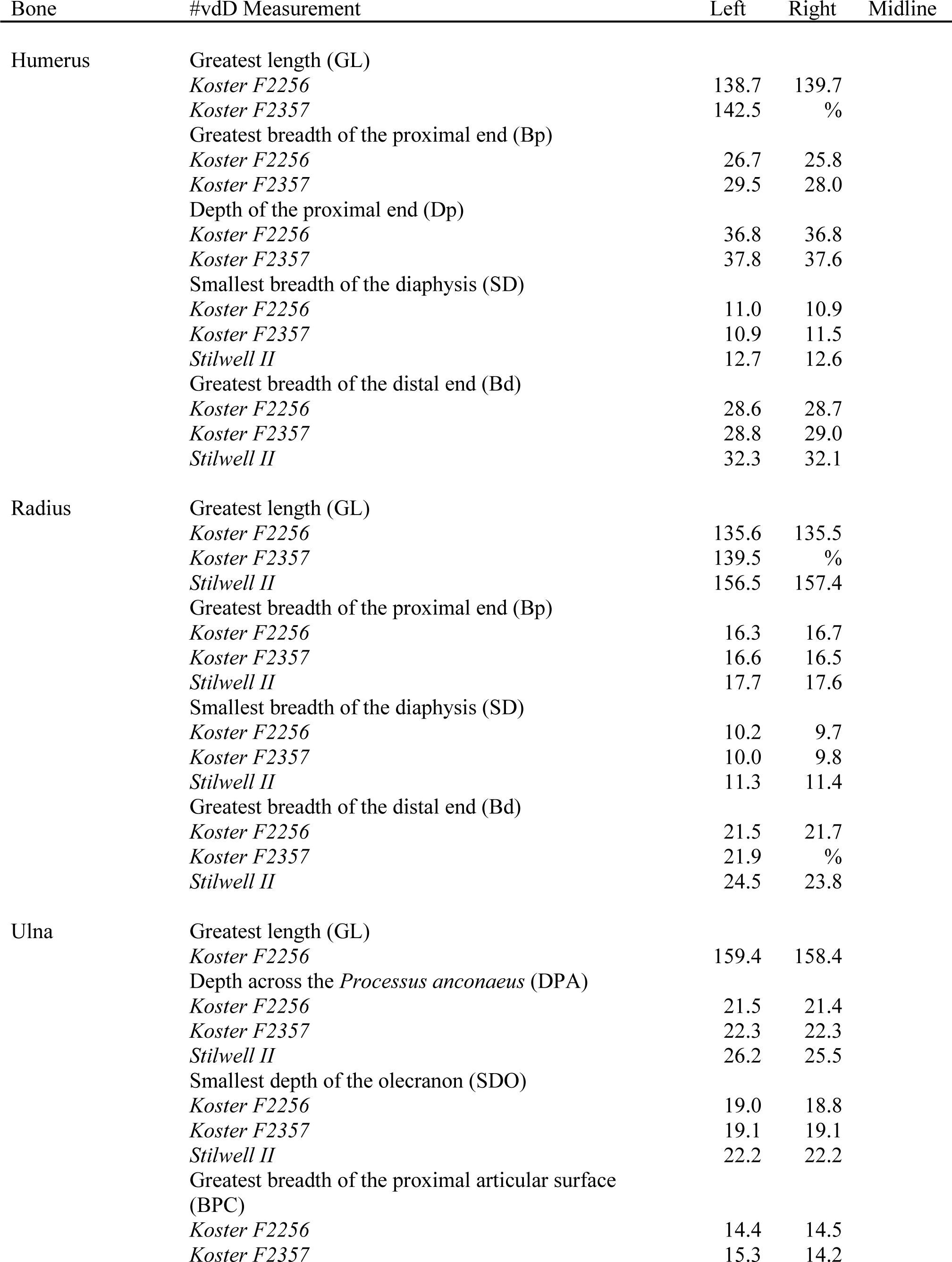
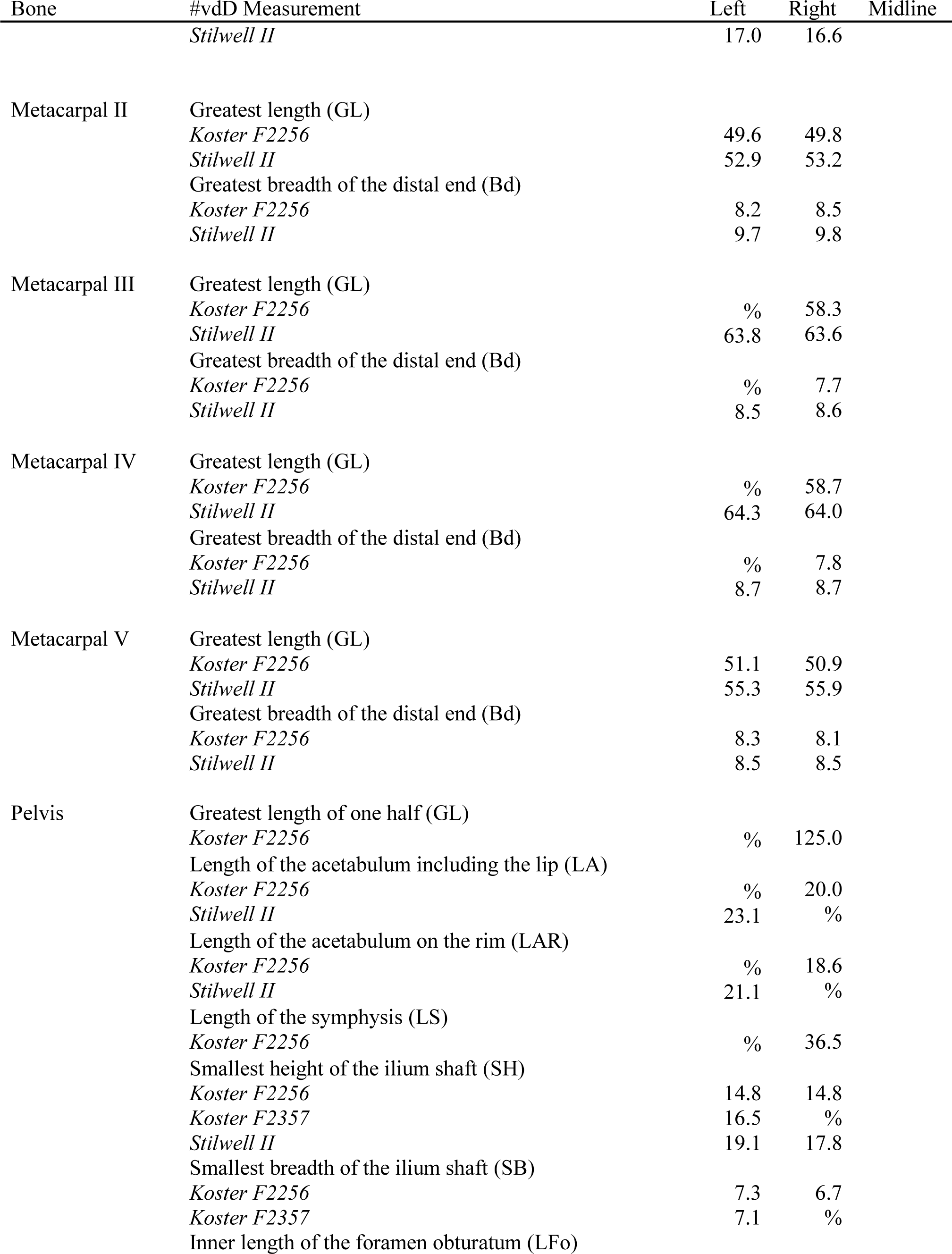
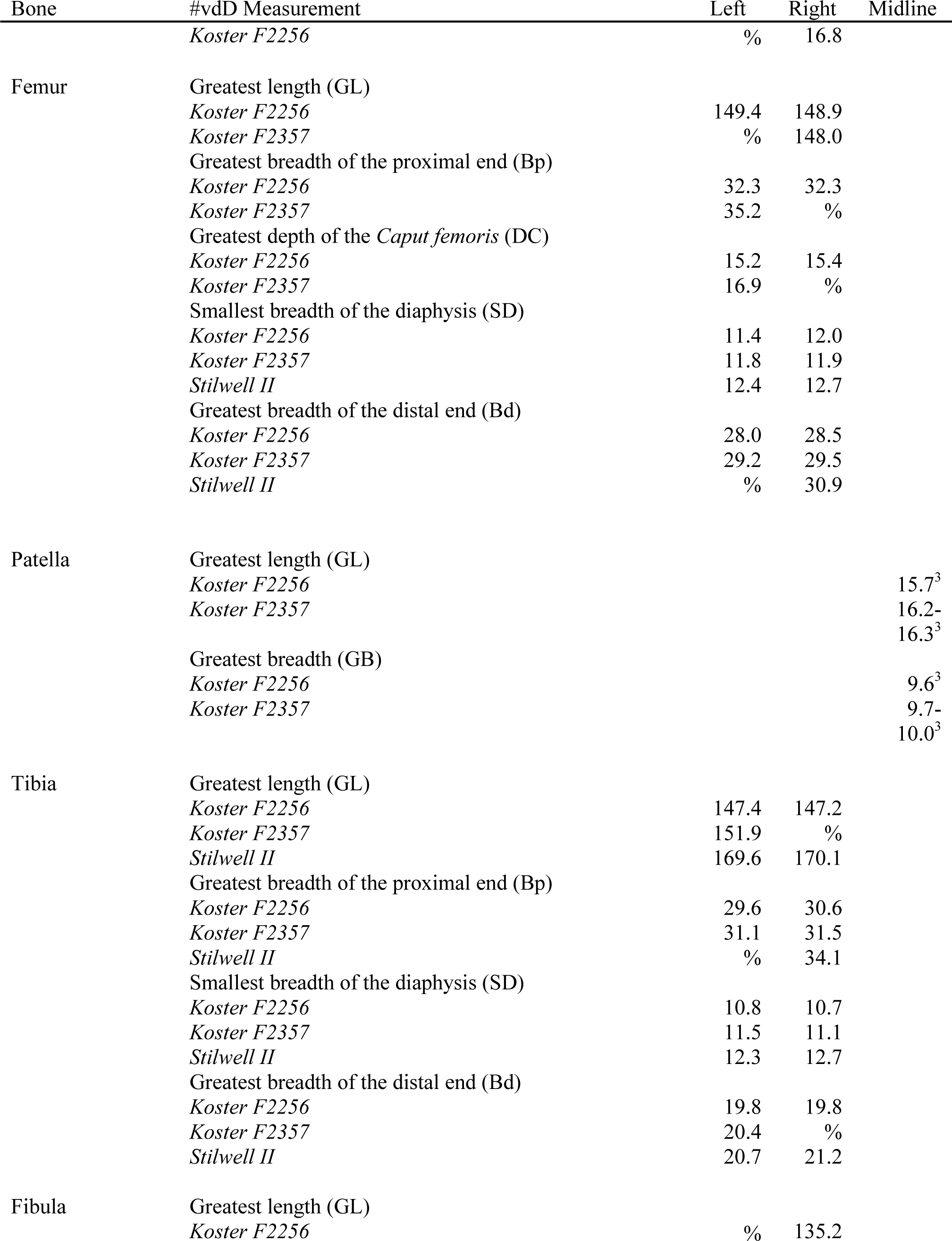
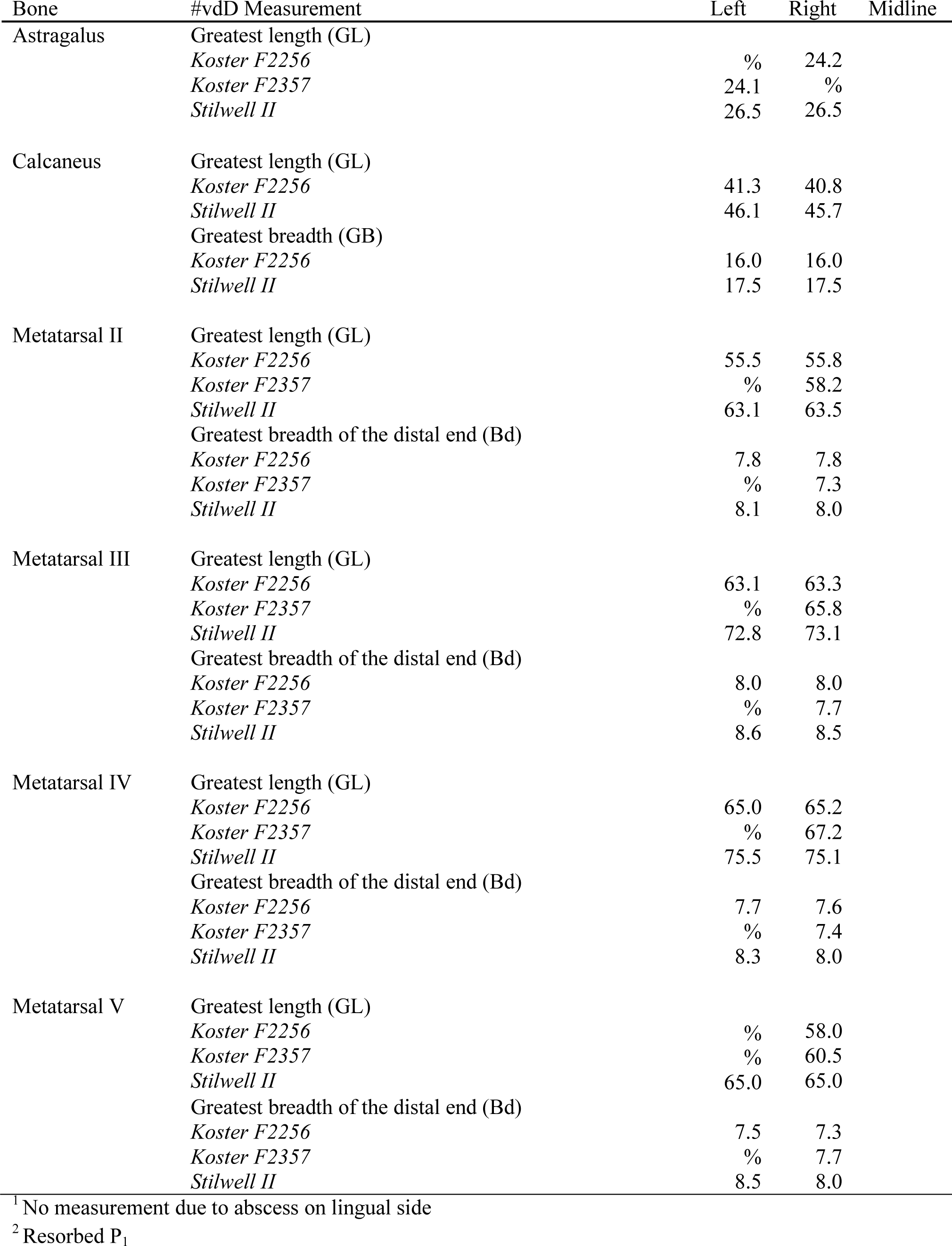
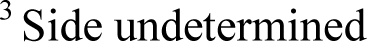
Measurements (mm) of the Stilwell II and Koster dogs, following von den Driesch (1976).

**Supplementary Table 2.**
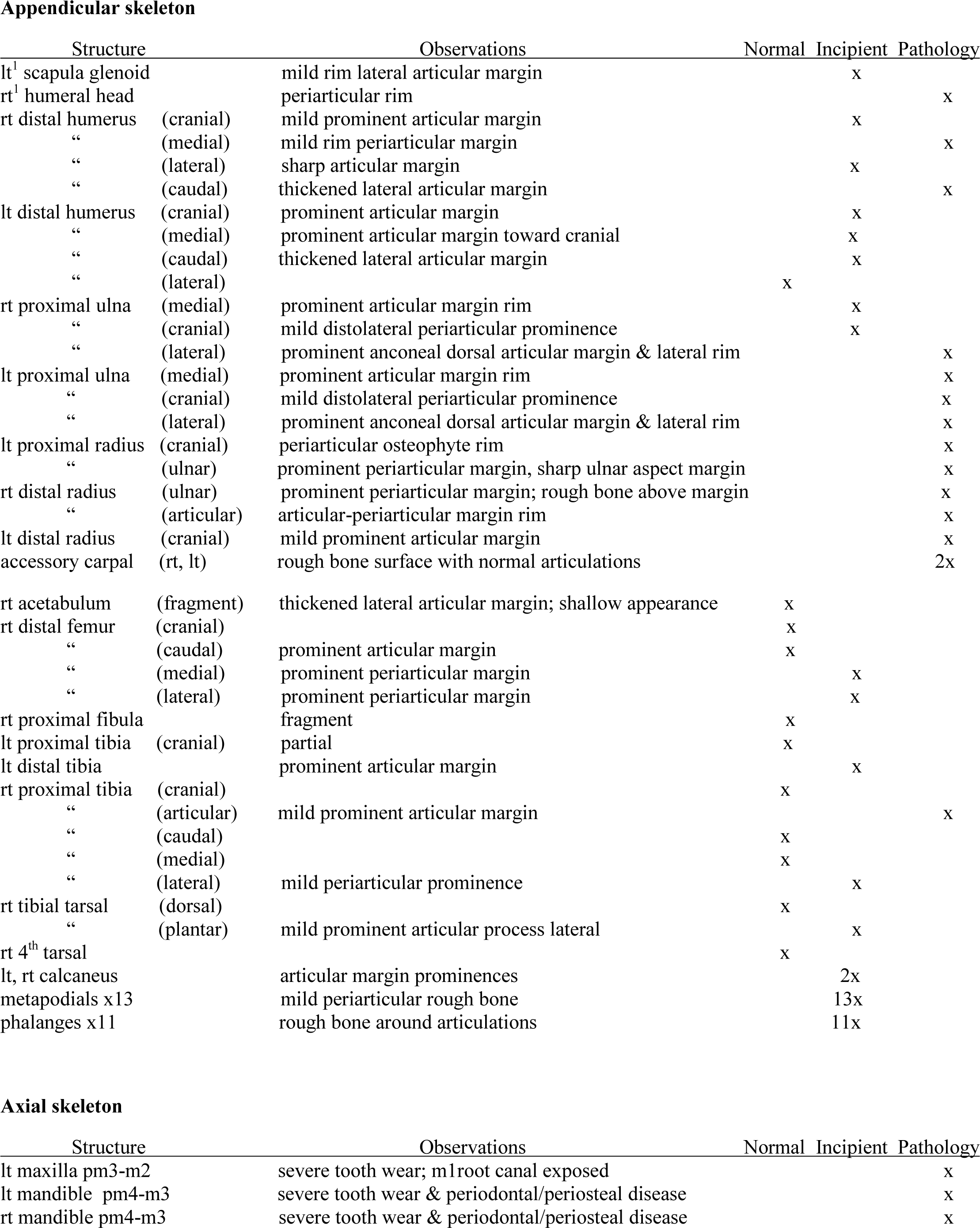
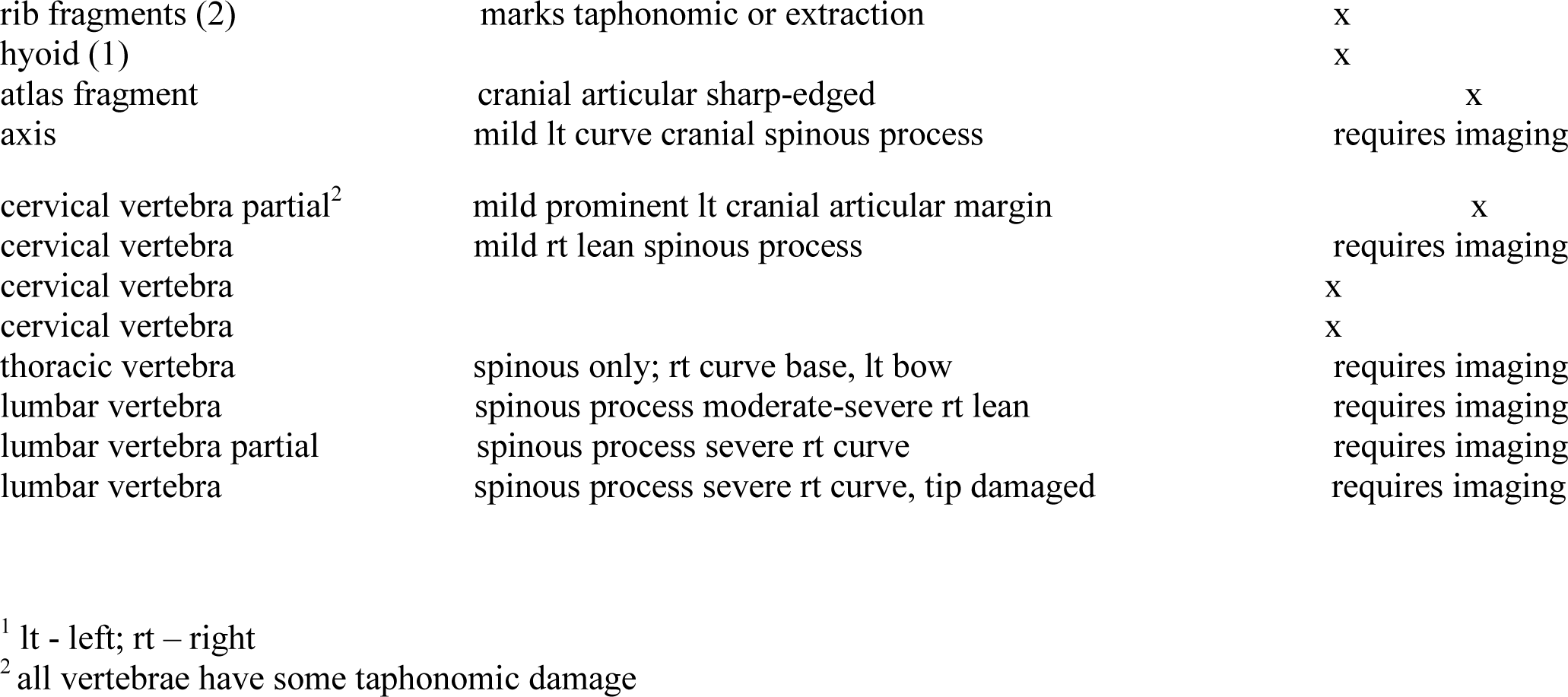
Skeletal Pathology Observations of the Stilwell II Dog

**Supplementary Table 3.**
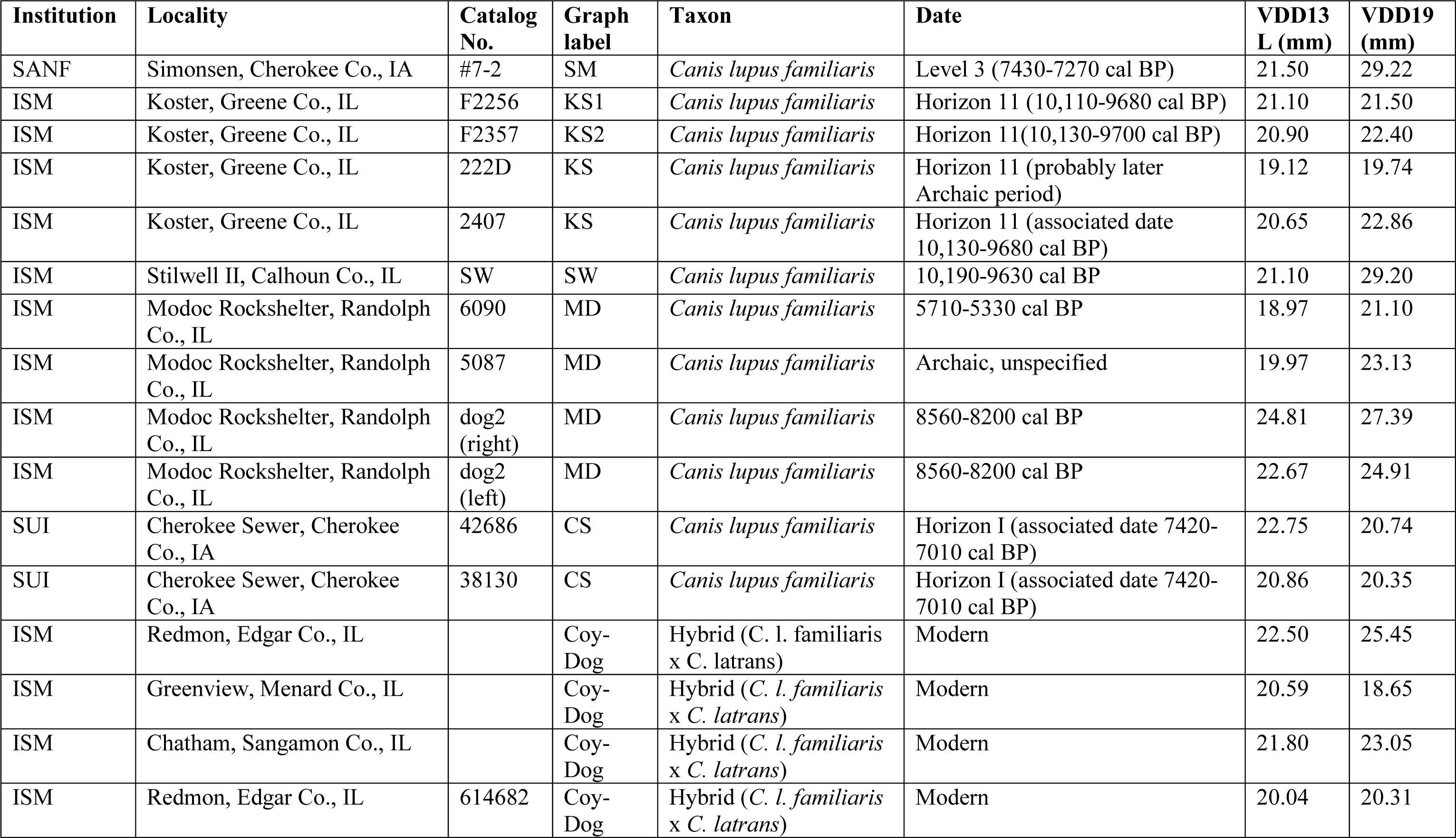
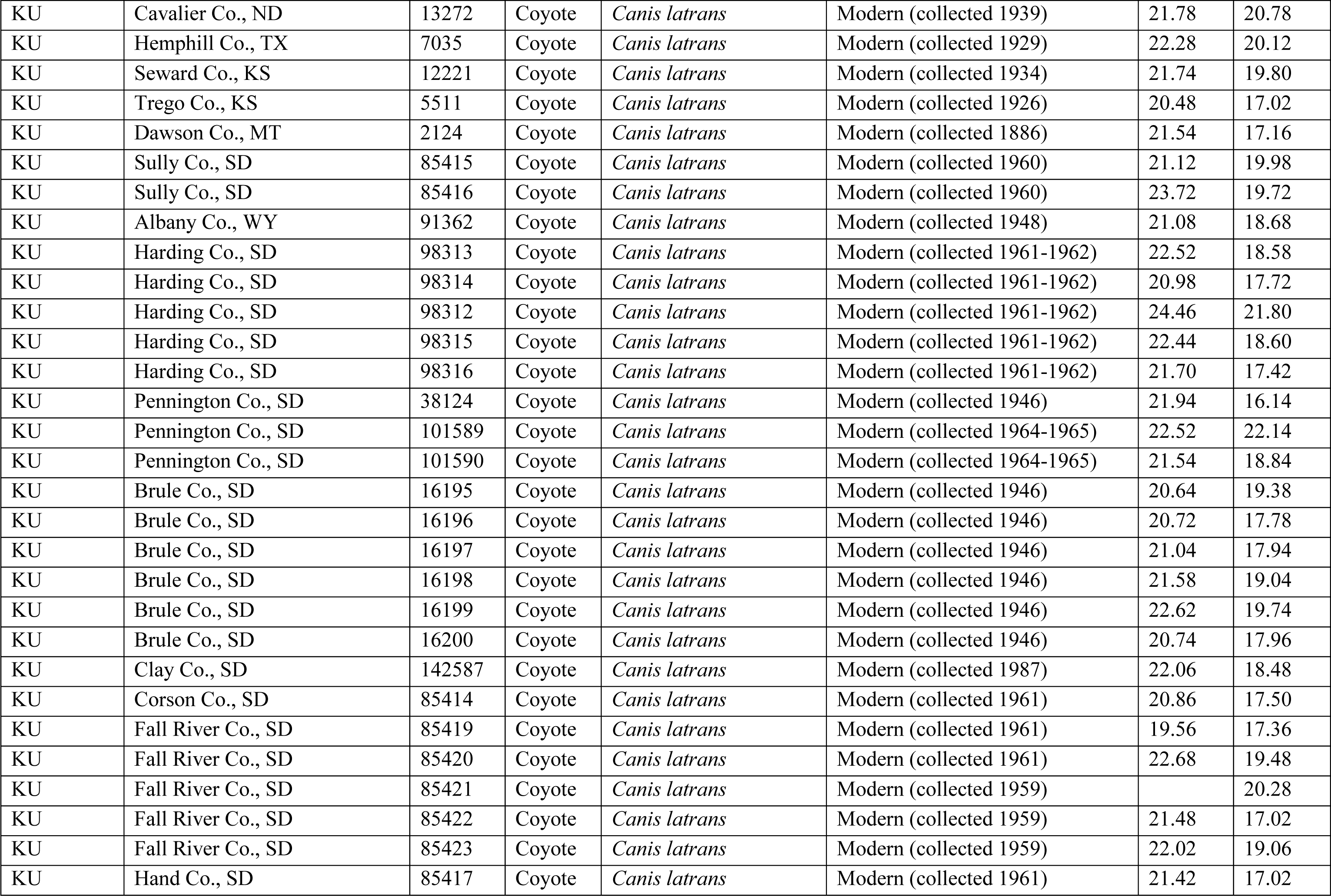
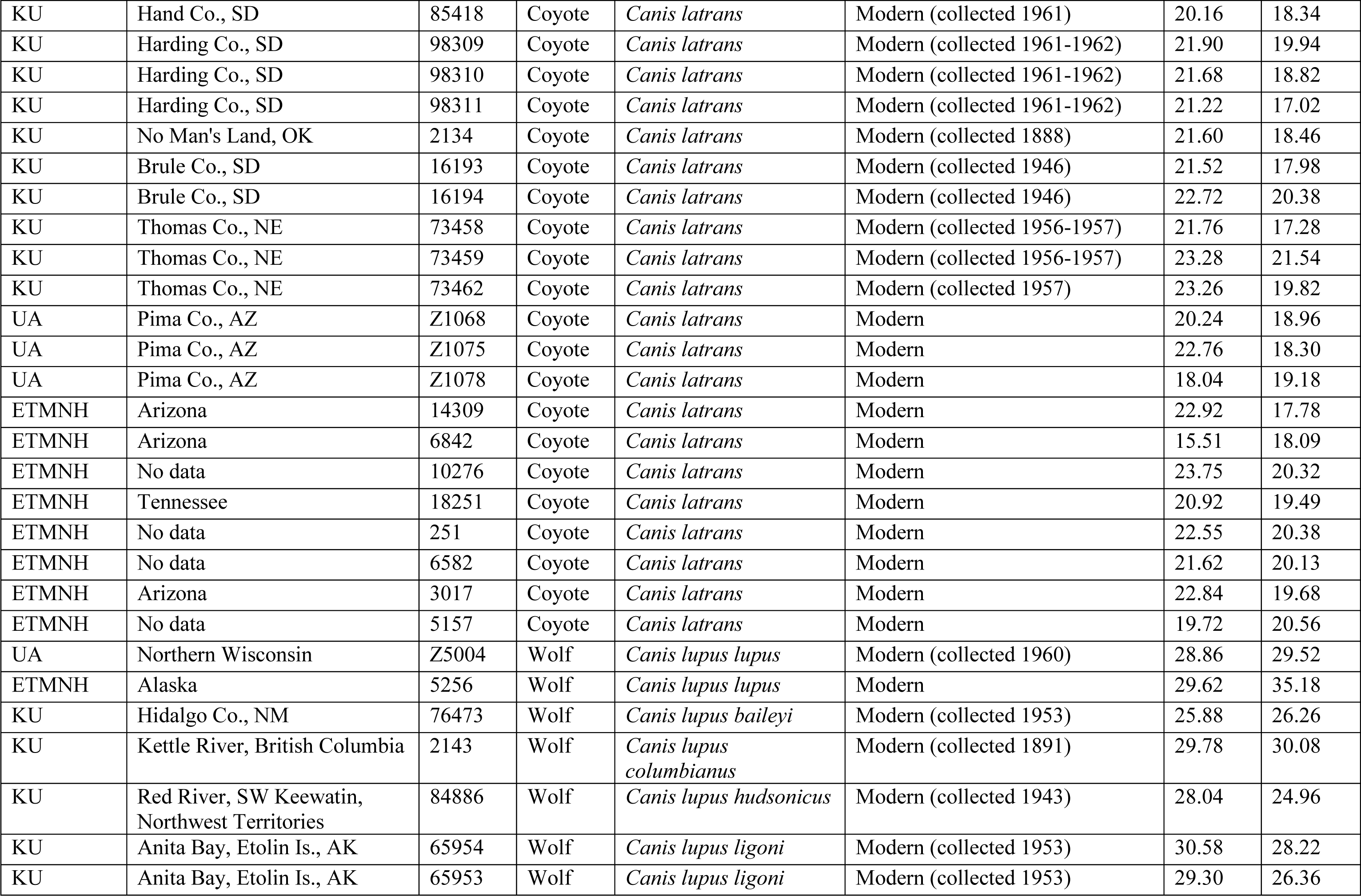
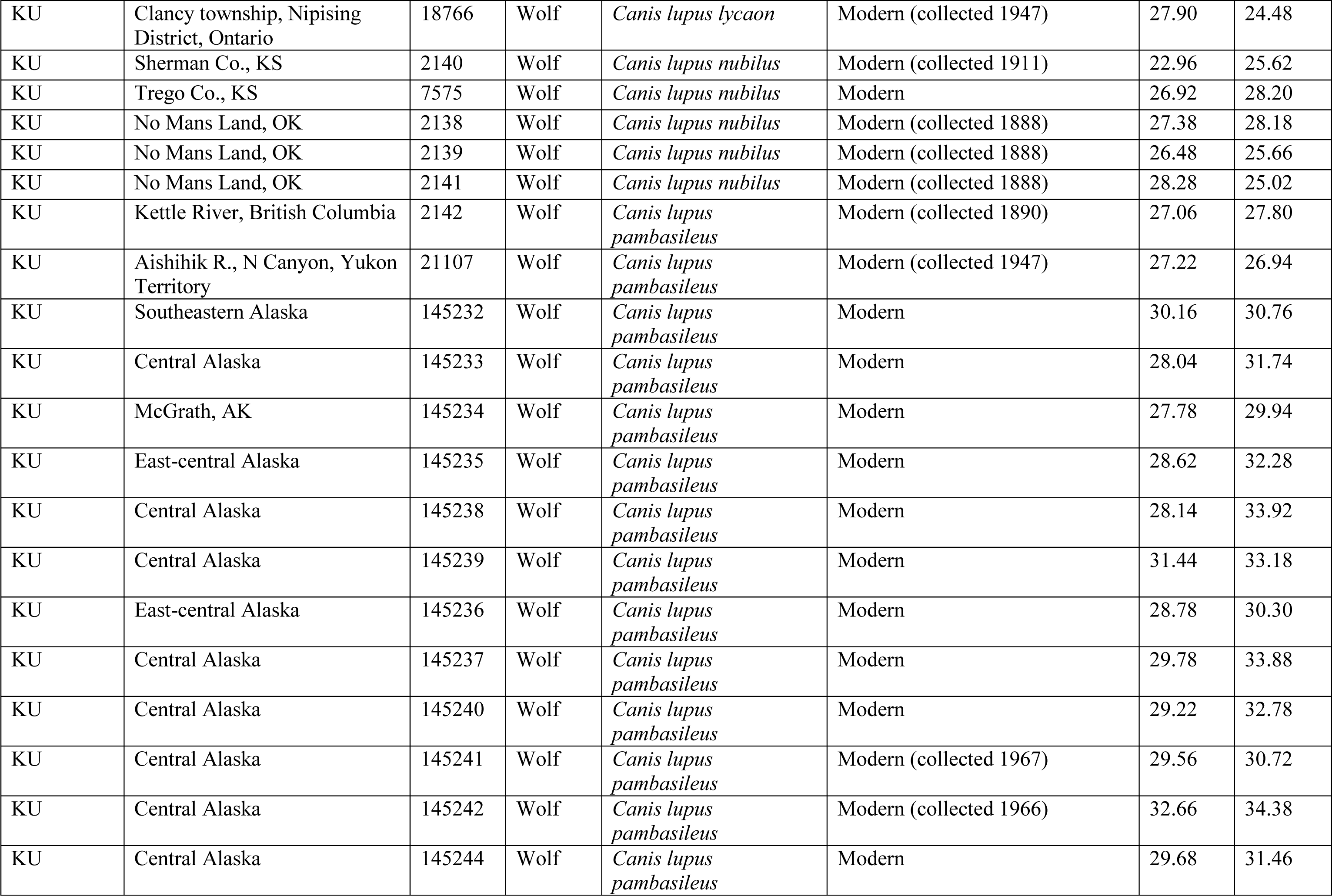
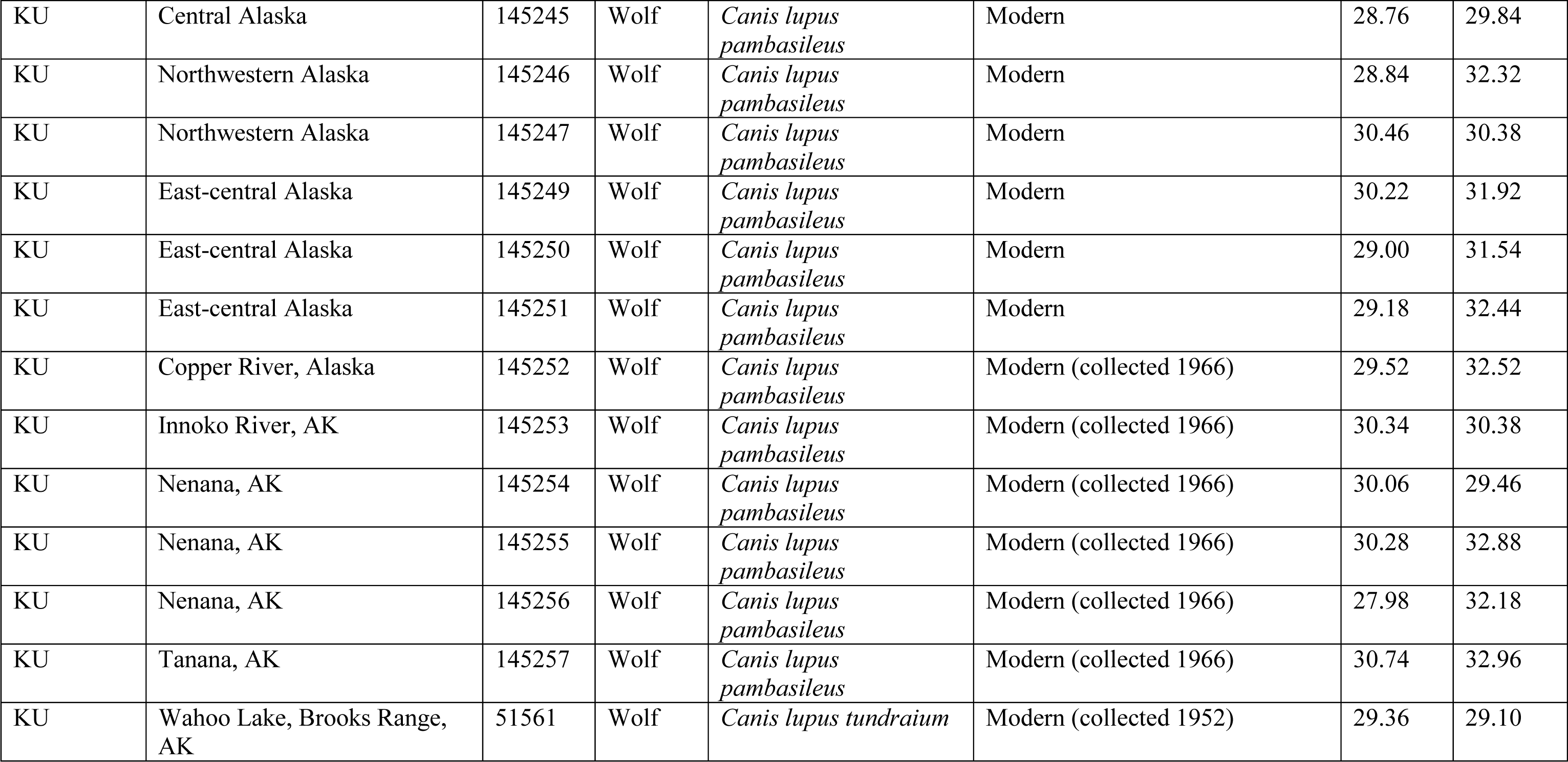
Comparative measurements on “length of camassial Mi” (VDD13L) and “height of mandible behind Mi” (VDD19), following von den Driesch (1976). Abbreviations as follows, SANF=Sanford Museum and Planetarium, Cherokee, IA; ISM=Illinois State Museum, Springfield, IL; KU=University of Kansas Natural History Museum, Lawrence, KS; UA=Stanley J. Olsen Laboratory of Zooarchaeology, Arizona State Museum, Tucson, AZ; ETMNH=East Tennessee State University Museum of Natural History, Johnson City, TN.

| Institution | Locality | Catalog No. | Graph label | Taxon | Date | VDD13 L (mm) | VDD19 (mm) |
| --- | --- | --- | --- | --- | --- | --- | --- |
| SANF | Simonsen, Cherokee Co., IA | #7-2 | SM | *Canis lupus familiaris* | Level 3 (7430-7270 cal BP) | 21.50 | 29.22 |
| ISM | Koster, Greene Co., IL | F2256 | KS1 | *Canis lupus familiaris* | Horizon 11 (10,110-9680 cal BP) | 21.10 | 21.50 |
| ISM | Koster, Greene Co., IL | F2357 | KS2 | *Canis lupus familiaris* | Horizon 11(10,130-9700 cal BP) | 20.90 | 22.40 |
| ISM | Koster, Greene Co., IL | 222D | KS | *Canis lupus familiaris* | Horizon 11 (probably later Archaic period) | 19.12 | 19.74 |
| ISM | Koster, Greene Co., IL | 2407 | KS | *Canis lupus familiaris* | Horizon 11 (associated date 10,130-9680 cal BP) | 20.65 | 22.86 |
| ISM | Stilwell II, Calhoun Co., IL | SW | SW | *Canis lupus familiaris* | 10,190-9630 cal BP | 21.10 | 29.20 |
| ISM | Modoc Rockshelter, Randolph Co., IL | 6090 | MD | *Canis lupus familiaris* | 5710-5330 cal BP | 18.97 | 21.10 |
| ISM | Modoc Rockshelter, Randolph Co., IL | 5087 | MD | *Canis lupus familiaris* | Archaic, unspecified | 19.97 | 23.13 |
| ISM | Modoc Rockshelter, Randolph Co., IL | dog2 (right) | MD | *Canis lupus familiaris* | 8560-8200 cal BP | 24.81 | 27.39 |
| ISM | Modoc Rockshelter, Randolph Co., IL | dog2 (left) | MD | *Canis lupus familiaris* | 8560-8200 cal BP | 22.67 | 24.91 |
| SUI | Cherokee Sewer, Cherokee Co., IA | 42686 | CS | *Canis lupus familiaris* | Horizon I (associated date 7420-7010 cal BP) | 22.75 | 20.74 |
| SUI | Cherokee Sewer, Cherokee Co., IA | 38130 | CS | *Canis lupus familiaris* | Horizon I (associated date 7420-7010 cal BP) | 20.86 | 20.35 |
| ISM | Redmon, Edgar Co., IL |  | Coy-Dog | Hybrid (*C. l. familiaris* x *C. latrans*) | Modern | 22.50 | 25.45 |
| ISM | Greenvew, Menard Co., IL |  | Coy-Dog | Hybrid (*C. l. familiaris* x *C. latrans*) | Modern | 20.59 | 18.65 |
| ISM | Chatham, Sangamon Co., IL |  | Coy-Dog | Hybrid (*C. l. familiaris* x *C. latrans*) | Modern | 21.80 | 23.05 |
| ISM | Redmon, Edgar Co., IL | 614682 | Coy-Dog | Hybrid (*C. l. familiaris* x *C. latrans*) | Modern | 20.04 | 20.31 |
| KU | Cavalier Co., ND | 13272 | Coyote | *Canis latrans* | Modern (collected 1939) | 21.78 | 20.78 |
| KU | Hemphill Co., TX | 7035 | Coyote | *Canis latrans* | Modern (collected 1929) | 22.28 | 20.12 |
| KU | Seward Co., KS | 12221 | Coyote | *Canis latrans* | Modern (collected 1934) | 21.74 | 19.80 |
| KU | Trego Co., KS | 5511 | Coyote | *Canis latrans* | Modern (collected 1926) | 20.48 | 17.02 |
| KU | Dawson Co., MT | 2124 | Coyote | *Canis latrans* | Modern (collected 1886) | 21.54 | 17.16 |
| KU | Sully Co., SD | 85415 | Coyote | *Canis latrans* | Modern (collected 1960) | 21.12 | 19.98 |
| KU | Sully Co., SD | 85416 | Coyote | *Canis latrans* | Modern (collected 1960) | 23.72 | 19.72 |
| KU | Albany Co., WY | 91362 | Coyote | *Canis latrans* | Modern (collected 1948) | 21.08 | 18.68 |
| KU | Harding Co., SD | 98313 | Coyote | *Canis latrans* | Modern (collected 1961-1962) | 22.52 | 18.58 |
| KU | Harding Co., SD | 98314 | Coyote | *Canis latrans* | Modern (collected 1961-1962) | 20.98 | 17.72 |
| KU | Harding Co., SD | 98312 | Coyote | *Canis latrans* | Modern (collected 1961-1962) | 24.46 | 21.80 |
| KU | Harding Co., SD | 98315 | Coyote | *Canis latrans* | Modern (collected 1961-1962) | 22.44 | 18.60 |
| KU | Harding Co., SD | 98316 | Coyote | *Canis latrans* | Modern (collected 1961-1962) | 21.70 | 17.42 |
| KU | Pennington Co., SD | 38124 | Coyote | *Canis latrans* | Modern (collected 1946) | 21.94 | 16.14 |
| KU | Pennington Co., SD | 101589 | Coyote | *Canis latrans* | Modern (collected 1964-1965) | 22.52 | 22.14 |
| KU | Pennington Co., SD | 101590 | Coyote | *Canis latrans* | Modern (collected 1964-1965) | 21.54 | 18.84 |
| KU | Brule Co., SD | 16195 | Coyote | *Canis latrans* | Modern (collected 1946) | 20.64 | 19.38 |
| KU | Brule Co., SD | 16196 | Coyote | *Canis latrans* | Modern (collected 1946) | 20.72 | 17.78 |
| KU | Brule Co., SD | 16197 | Coyote | *Canis latrans* | Modern (collected 1946) | 21.04 | 17.94 |
| KU | Brule Co., SD | 16198 | Coyote | *Canis latrans* | Modern (collected 1946) | 21.58 | 19.04 |
| KU | Brule Co., SD | 16199 | Coyote | *Canis latrans* | Modern (collected 1946) | 22.62 | 19.74 |
| KU | Brule Co., SD | 16200 | Coyote | *Canis latrans* | Modern (collected 1946) | 20.74 | 17.96 |
| KU | Clay Co., SD | 142587 | Coyote | *Canis latrans* | Modern (collected 1987) | 22.06 | 18.48 |
| KU | Corson Co., SD | 85414 | Coyote | *Canis latrans* | Modern (collected 1961) | 20.86 | 17.50 |
| KU | Fall River Co., SD | 85419 | Coyote | *Canis latrans* | Modern (collected 1961) | 19.56 | 17.36 |
| KU | Fall River Co., SD | 85420 | Coyote | *Canis latrans* | Modern (collected 1961) | 22.68 | 19.48 |
| KU | Fall River Co., SD | 85421 | Coyote | *Canis latrans* | Modern (collected 1959) |  | 20.28 |
| KU | Fall River Co., SD | 85422 | Coyote | *Canis latrans* | Modern (collected 1959) | 21.48 | 17.02 |
| KU | Fall River Co., SD | 85423 | Coyote | *Canis latrans* | Modern (collected 1959) | 22.02 | 19.06 |
| KU | Hand Co., SD | 85417 | Coyote | *Canis latrans* | Modern (collected 1961) | 21.42 | 17.02 |
| KU | Hand Co., SD | 85418 | Coyote | *Canis latrans* | Modern (collected 1961) | 20.16 | 18.34 |
| KU | Harding Co., SD | 98309 | Coyote | *Canis latrans* | Modern (collected 1961-1962) | 21.90 | 19.94 |
| KU | Harding Co., SD | 98310 | Coyote | *Canis latrans* | Modern (collected 1961-1962) | 21.68 | 18.82 |
| KU | Harding Co., SD | 98311 | Coyote | *Canis latrans* | Modern (collected 1961-1962) | 21.22 | 17.02 |
| KU | No Man's Land, OK | 2134 | Coyote | *Canis latrans* | Modern (collected 1888) | 21.60 | 18.46 |
| KU | Brule Co., SD | 16193 | Coyote | *Canis latrans* | Modern (collected 1946) | 21.52 | 17.98 |
| KU | Brule Co., SD | 16194 | Coyote | *Canis latrans* | Modern (collected 1946) | 22.72 | 20.38 |
| KU | Thomas Co., NE | 73458 | Coyote | *Canis latrans* | Modern (collected 1956-1957) | 21.76 | 17.28 |
| KU | Thomas Co., NE | 73459 | Coyote | *Canis latrans* | Modern (collected 1956-1957) | 23.28 | 21.54 |
| KU | Thomas Co., NE | 73462 | Coyote | *Canis latrans* | Modern (collected 1957) | 23.26 | 19.82 |
| UA | Pima Co., AZ | Z1068 | Coyote | *Canis latrans* | Modern | 20.24 | 18.96 |
| UA | Pima Co., AZ | Z1075 | Coyote | *Canis latrans* | Modern | 22.76 | 18.30 |
| UA | Pima Co., AZ | Z1078 | Coyote | *Canis latrans* | Modern | 18.04 | 19.18 |
| ETMNH | Arizona | 14309 | Coyote | *Canis latrans* | Modern | 22.92 | 17.78 |
| ETMNH | Arizona | 6842 | Coyote | *Canis latrans* | Modern | 15.51 | 18.09 |
| ETMNH | No data | 10276 | Coyote | *Canis latrans* | Modern | 23.75 | 20.32 |
| ETMNH | Tennessee | 18251 | Coyote | *Canis latrans* | Modern | 20.92 | 19.49 |
| ETMNH | No data | 251 | Coyote | *Canis latrans* | Modern | 22.55 | 20.38 |
| ETMNH | No data | 6582 | Coyote | *Canis latrans* | Modern | 21.62 | 20.13 |
| ETMNH | Arizona | 3017 | Coyote | *Canis latrans* | Modern | 22.84 | 19.68 |
| ETMNH | No data | 5157 | Coyote | *Canis latrans* | Modern | 19.72 | 20.56 |
| UA | Northern Wisconsin | Z5004 | Wolf | *Canis lupus lupus* | Modern (collected 1960) | 28.86 | 29.52 |
| ETMNH | Alaska | 5256 | Wolf | *Canis lupus lupus* | Modern | 29.62 | 35.18 |
| KU | Hidalgo Co., NM | 76473 | Wolf | *Canis lupus baileyi* | Modern (collected 1953) | 25.88 | 26.26 |
| KU | Kettle River, British Columbia | 2143 | Wolf | *Canis lupus columbianus* | Modern (collected 1891) | 29.78 | 30.08 |
| KU | Red River, SW Keewatin, Northwest Territories | 84886 | Wolf | *Canis lupus hudsonicus* | Modern (collected 1943) | 28.04 | 24.96 |
| KU | Anita Bay, Etolin Is., AK | 65954 | Wolf | *Canis lupus ligoni* | Modern (collected 1953) | 30.58 | 28.22 |
| KU | Anita Bay, Etolin Is., AK | 65953 | Wolf | *Canis lupus ligoni* | Modern (collected 1953) | 29.30 | 26.36 |
| KU | Clancy township, Nipising District, Ontario | 18766 | Wolf | *Canis lupus lycaon* | Modern (collected 1947) | 27.90 | 24.48 |
| KU | Sherman Co., KS | 2140 | Wolf | *Canis lupus nubilus* | Modern (collected 1911) | 22.96 | 25.62 |
| KU | Trego Co., KS | 7575 | Wolf | *Canis lupus nubilus* | Modern | 26.92 | 28.20 |
| KU | No Mans Land, OK | 2138 | Wolf | *Canis lupus nubilus* | Modern (collected 1888) | 27.38 | 28.18 |
| KU | No Mans Land, OK | 2139 | Wolf | *Canis lupus nubilus* | Modern (collected 1888) | 26.48 | 25.66 |
| KU | No Mans Land, OK | 2141 | Wolf | *Canis lupus nubilus* | Modern (collected 1888) | 28.28 | 25.02 |
| KU | Kettle River, British Columbia | 2142 | Wolf | *Canis lupus pambasileus* | Modern (collected 1890) | 27.06 | 27.80 |
| KU | Aishihik R., N Canyon, Yukon Territory | 21107 | Wolf | *Canis lupus pambasileus* | Modern (collected 1947) | 27.22 | 26.94 |
| KU | Southeastern Alaska | 145232 | Wolf | *Canis lupus pambasileus* | Modern | 30.16 | 30.76 |
| KU | Central Alaska | 145233 | Wolf | *Canis lupus pambasileus* | Modern | 28.04 | 31.74 |
| KU | McGrath, AK | 145234 | Wolf | *Canis lupus pambasileus* | Modern | 27.78 | 29.94 |
| KU | East-central Alaska | 145235 | Wolf | *Canis lupus pambasileus* | Modern | 28.62 | 32.28 |
| KU | Central Alaska | 145238 | Wolf | *Canis lupus pambasileus* | Modern | 28.14 | 33.92 |
| KU | Central Alaska | 145239 | Wolf | *Canis lupus pambasileus* | Modern | 31.44 | 33.18 |
| KU | East-central Alaska | 145236 | Wolf | *Canis lupus pambasileus* | Modern | 28.78 | 30.30 |
| KU | Central Alaska | 145237 | Wolf | *Canis lupus pambasileus* | Modern | 29.78 | 33.88 |
| KU | Central Alaska | 145240 | Wolf | *Canis lupus pambasileus* | Modern | 29.22 | 32.78 |
| KU | Central Alaska | 145241 | Wolf | *Canis lupus pambasileus* | Modern (collected 1967) | 29.56 | 30.72 |
| KU | Central Alaska | 145242 | Wolf | *Canis lupus pambasileus* | Modern (collected 1966) | 32.66 | 34.38 |
| KU | Central Alaska | 145244 | Wolf | *Canis lupus pambasileus* | Modern | 29.68 | 31.46 |
| KU | Central Alaska | 145245 | Wolf | *Canis lupus pambasileus* | Modern | 28.76 | 29.84 |
| KU | Northwestern Alaska | 145246 | Wolf | *Canis lupus pambasileus* | Modern | 28.84 | 32.32 |
| KU | Northwestern Alaska | 145247 | Wolf | *Canis lupus pambasileus* | Modern | 30.46 | 30.38 |
| KU | East-central Alaska | 145249 | Wolf | *Canis lupus pambasileus* | Modern | 30.22 | 31.92 |
| KU | East-central Alaska | 145250 | Wolf | *Canis lupus pambasileus* | Modern | 29.00 | 31.54 |
| KU | East-central Alaska | 145251 | Wolf | *Canis lupus pambasileus* | Modern | 29.18 | 32.44 |
| KU | Copper River, Alaska | 145252 | Wolf | *Canis lupus pambasileus* | Modern (collected 1966) | 29.52 | 32.52 |
| KU | Innoko River, AK | 145253 | Wolf | *Canis lupus pambasileus* | Modern (collected 1966) | 30.34 | 30.38 |
| KU | Nenana, AK | 145254 | Wolf | *Canis lupus pambasileus* | Modern (collected 1966) | 30.06 | 29.46 |
| KU | Nenana, AK | 145255 | Wolf | *Canis lupus pambasileus* | Modern (collected 1966) | 30.28 | 32.88 |
| KU | Nenana, AK | 145256 | Wolf | *Canis lupus pambasileus* | Modern (collected 1966) | 27.98 | 32.18 |
| KU | Tanana, AK | 145257 | Wolf | *Canis lupus pambasileus* | Modern (collected 1966) | 30.74 | 32.96 |
| KU | Wahoo Lake, Brooks Range, AK | 51561 | Wolf | *Canis lupus tundraiium* | Modern (collected 1952) | 29.36 | 29.10 |

